# Out of the blue: Phototropins of the leaf vascular bundle sheath mediate the regulation of leaf hydraulic conductance by blue light

**DOI:** 10.1101/720912

**Authors:** Yael Grunwald, Sanbon Chaka Gosa, Tanmayee Torne Srivastava, Nava Moran, Menachem Moshelion

## Abstract

The Arabidopsis leaf veins bundle sheath cells (BSCs) – a selective barrier to water and solutes entering the mesophyll – increase the leaf radial hydraulic conductance (K_leaf_) by acidifying the xylem sap by their plasma membrane H^+^-ATPase, AHA2. Based on this and on the BSCs’ expression of *PHOT1* and *PHOT2,* and the known blue-light (BL)-induced K_leaf_ increase, we hypothesized that, resembling the guard cells, BL perception by the BSCs’ phots activates its H^+^-ATPase, which, consequently, upregulates K_leaf_. Indeed, under BL, the K_leaf_ of the knockout mutant lines *phot1-5*, *phot2-1*, *phot1-5phot2-1,* and *aha2-4* was lower than that of the WT. BSC-only-directed complementation of *phot1-5* or *aha2-4* by *PHOT1* or *AHA2*, respectively, restored the BL-induced K_leaf_ increase. BSC-specific silencing of *PHOT1* or *PHOT2* prevented such K_leaf_ increase. A xylem-fed kinase inhibitor (tyrphostin 9) replicated this also in WT plants. White light – ineffective in the *phot1-5* mutant – acidified the xylem sap (relative to darkness) in WT and in the *PHOT1*-complemented *phot1-5*. These results, supported by BL increase of BSC protoplasts’ water permeability and cytosolic pH and their hyperpolarization by BL, identify the BSCs as a second phot-controlled water conductance element in leaves, in series with stomatal conductance. Through both, BL regulates the leaf water balance.

**One-Sentence summary:** Blue light regulates the leaf hydraulic conductance via the bundle-sheath cells’ blue light PHOT receptors which, via an autonomous signaling pathway, activate the BSCs’ AHA2 H^+^-pump.

## Introduction

Light, the energy source for photosynthesis, also regulates plant growth and development (Kronenberg and Kendrick, 1986) as well as physiological traits such as stomatal conductance (g_s_; Zeiger and Helper, 1977; Karlsson, 1986; Talbott et al., 2003) and leaf hydraulic conductance (K_leaf_; Voicu et al., 2008; Ben Baaziz et al., 2012; Aasamaa and Sõber, 2012; Prado and Maurel, 2013). One of the most conserved and well-studied light-sensing mechanisms is the sensing of blue light (BL, 390–550 nm) that causes stomata to open (Grondin et al., 2015). In this signal transduction pathway in the guard cells (GCs), BL is perceived by the protein kinases phot1 and phot2 (Kinoshita et al., 2001). Their immediate phosphorylation substrate is BLUS1 (Blue Light Signaling1, At4g14480; Takemiya et al., 2013), which is a part of the earliest stomata-opening BL signaling complex with phot. Another part of this complex is the Raf-like kinase, BHP (BL-dependent H^+^-ATPase phosphorylation, At4g18950; Hayashi et al., 2017; reviewed by Inoue and Kinoshita, 2017). While none of these directly phosphorylates the guard-cell H^+^-ATPase (Hayashi et al., 2017), one of the other Raf-like kinases downstream of phot could be the target of tyrphostin 9, a kinase inhibitor that has been shown to inhibit the BL-induced phosphorylation of the penultimate threonine of the plasma membrane H^+^-ATPase (AHA) in Arabidopsis GCs (ibid). This BL-induced phosphorylation activates the AHA (Kinoshita and Shimazaki, 1999; Svennelid et al., 1999), which, in turn, hyperpolarizes the GCs and acidifies their surrounding apoplast (Kinoshita and Shimazaki, 1999; Ueno et al., 2005; Den Os et al., 2007; Elmore and Coaker, 2011, reviewed by Inoue and Kinoshita, 2017). Conversely, at night, in the dark, the guard cells’ H^+^-ATPases are deactivated, the GCs are depolarized, and their apoplast becomes alkaline (Kinoshita et al., 2001; Kinoshita and Shimazaki, 1999). These pH and membrane potential changes cause ion and water fluxes across the GCs membranes causing their osmotic swelling during the day and shrinking at night, which underlies stomata opening and closure (Assmann, 1993; Roelfsema and Hedrich, 2005; Shimazaki et al., 2007).

Both BL and white light (WL) have been shown to increase the hydraulic conductance (K_leaf_) of the entire leaf in several plant species [(Voicu et al., 2008 (bur oak, *Quercus macrocarpa*); Voicu et al., 2009; Sellin et al., 2011 (silver birch, *Betula pendula* Roth); Aasamaa and Sber, 2012 (deciduous trees); Ben Baaziz et al., 2012 (walnut, *Juglans regia* L.)]. Interestingly, K_leaf_ responds more quickly to light than g_S_ does (Guyot et al., 2012). However, the molecular mechanism by which K_leaf_ increases in response to light quality is not yet fully understood.

In the past decade, it has been established that bundle sheath cells (BSCs), which form a parenchymal layer that tightly enwraps the entire leaf vasculature, can act as a dynamic and selective xylem-mesophyll barrier to water and ions, and participate in K_leaf_ regulation (Shapira et al., 2009; Shatil-Cohen and Moshelion, 2012). Moreover, we recently discovered that the BSCs’ H^+^-ATPase proton pump, AHA2, participates in regulating K_leaf_ via changes in the xylem pH and that the positive correlation observed between AHA2-driven xylem acidification and K_leaf_ is due to an increase in the osmotic water permeability of BSCs’ membranes (Grunwald et al., 2021). AHA2 is particularly abundant in BSCs (present at a >3-fold higher level than that seen in neighboring mesophyll cells; Wigoda et al., 2017) which explains how such acidification is possible. In addition, the same BSC transcriptome analysis (ibid.) revealed substantial expression of the BL receptors phot1 and phot2 in BSCs, as well as substantial expression of the kinases BHP and BLUS1. These findings led us to hypothesize that the vascular BSCs possess a BL signal transduction pathway that is similar to the one that opens stomata and that plays a role in the regulation of K_leaf_. In confirmation of this hypothesis, we describe here a few of the physiological and molecular mechanisms underlying the blue light-dependent regulation of K_leaf_ by BSCs.

## Materials and Methods

### Plant material

#### Plant genotypes

We used WT (wild type) *Arabidopsis thaliana* plants ecotype Colombia, Col-0 (aka Col), the T-DNA insertion AHA2 mutants *aha2-4* (SALK_ 082786) (Col) and *aha2-4* complemented with *SCR: AHA2* (line 56) (Col), and *SCR:GFP* (Col) as described previously (Grunwald et al., 2021). In addition, we used WT Arabidopsis (Col) with the glabrous mutation (WT Col-gl), *phot1-5* (*nph1-5*) (Col-gl), *phot 2-1* (*npl1-1* or *cav1-1*) (Col-gl) and the double mutant *phot1-5phot2-1* (*nph1-5 npl1-1*) (Col-gl) (Kagawa et al., 2001), which were kindly provided by the Ken-Ichiro Shimazaki lab (Tokyo, Japan). The *blus1-4* (Col) seed was a kind gift from the Takemiya lab (Yamaguchi, Japan).

#### Construction of SCR:amiR-PHOT1 plants

The premiR-*PHOT1*or *PHOT2* and synthetic genes were synthesized by Hylabs (Rehovot, Israel), based on a premiR164 backbone (Alvarez et al., 2006). We used the Web-based MicroRNA Designer (WMD, http://wmd3.weigelworld.org) to produce a premiRNA gene MIR319a as described in WMD. After sequence verification, the premiR-*PHOT1*or premiR-*PHOT2* was cloned into the pDONR™ 221 and the SCR promoter into pDONRP4P1r (Invitrogen) vectors which are Gateway® compatible by BP reactions, and later cloned into the pB7M24GW (Invitrogen) two-fragment binary vector using an LR r reaction, according to the manufacturer’s instructions. Each binary vector was transformed into *Agrobacterium tumefaciens* by electroporation and transformed into WT Col-0 using the floral-dip method (Clough and Bent, 1998). Transformants were selected based on their resistance to BASTA herbicide (glufosinate ammonium, Sigma cat # 45520), grown on plates with MS (Murashige and Skoog, Duchefa cat# M222.0050) basal medium + 1 % sucrose and 20 µg/ml BASTA. DNA insertion was verified in selected lines by PCR targeting the junction of the premiR-gene and the 35S terminator with the forward primer located about 1000 bp from the 3’ end of premiR-gene and reverse primer on the 35S terminator (see primer list in Supplemental Table S1). PCR fragments were then sequenced and verified.

#### Construction of SCR: PHOT1 plants

Binary vectors were constructed with the *PHOT1* gene as described above and then transformed into *phot1-5* (Col-gl1) plants. Successful transformation was verified by PCR.

#### Plant growth conditions

Plants were grown in 250 ml pots, 2-3 plants per pot filled with soil (Klasmann686, Klasmann-Deilmann, Germany) + 4g/L Osmocote® 6M in a growth chamber kept at 22°C and 70% relative humidity, with an 8-h light (9:00 am–5:00 pm) /16-h dark photoperiod. During the day, the illumination, humidity and vapor pressure deficit (VPD) changed in a pattern resembling natural conditions, as in (Negin and Moshelion, 2017). The illumination intensity provided by LED lights strips [EnerLED 24 V-5630, 24 W/m, 3000 K (50%)/6000 K (50%)] reached 150–200 μmol m^−2^ sec^−1^ light at the plant level at midday. The plants were irrigated twice a week.

#### Preparation of detached leaves for gas-exchange measurements

Fully expanded leaves from 7- to 8-week-old plants were excised at the petiole base using a wet, sharp blade under a drop of water. Petioles were immediately submerged in 0.5-ml Eppendorf tubes with artificial xylem solution [AXS; 3 mM KNO_3_, 1 mM Ca(NO_3_)_2_, 1 mM MgSO_4_, 3 mM CaCl_2_, 0.25 mM NaH_2_PO_4_, 90 µM EDFS (Sigma)]. The leaves were excised, shortly before the “lights off” transition (around 5:00 pm) on the evening preceding the measurements and placed in gas-sealed, transparent 25 cm × 20 cm × 15 cm plastic boxes with water-soaked tissue paper on the bottom to provide ∼90% humidity. The transparent boxes were then inserted into a larger light-proof plastic box overnight and kept in total darkness until the start of light treatments the next morning.

### Light treatments of detached leaves

All gas-exchange experiments were conducted in a dark room (<1 umol m^-2^ s^-1^) at a constant temperature of 22°C, between 10:00 am and 1:00 pm. The excised leaves were taken out of the light-sealed box and placed in one of two custom-made light chambers. The total illumination intensity in both light chambers was set to roughly 220 µmol m^-2^ s^-1^. In the red-light chamber, the leaves were illuminated only with red light (RL, 660 nm) and in the RL+BL chamber they were illuminated with approximately 90% RL and 10% BL (450 nm). Leaves were exposed to either RL or RL+BL for 15 min at an ambient VPD of 1.3–1.5 kPa. A fan circulated the air in each light chamber for uniform and constant temperature and VPD. Next, leaves were transferred to a LI-COR 6400XT for an additional 2–3 min, under the same illumination conditions with VPD of about 1.3kPa, until stabilization of the gas-exchange measurements, after which a measurement was recorded. D-LED lighting with adjustable current source (DR-SD6/12) was used as light sources (http://www.d-led.net/). Light intensity was verified daily using a LI-250A light meter with a LI-190SA quantum sensor (LI-COR, Lincoln, NE, USA).

### Determination of gas exchange, E and g_S_, and hydraulic conductance, K_leaf_

Following the light treatments, the leaves were immediately placed in a LI-COR 6400XT for measurements of stomatal conductance (g_S_) and transpiration (E) similar to Grunwald et al. (2021), with the following changes: All of the experiments were conducted in the dark at 22°C and the illumination in the LI-COR cuvette was adjusted to match the preceding light treatments. Then, the water potential (Ψ_leaf_) was determined in a pressure chamber (ARIMAD-3000; MRC Israel) and K_leaf_ was calculated (as in Grunwald et al., 2021) as:

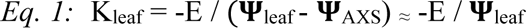

where AXS denotes artificial xylem sap (solution).

Additionally, g_S_ was measured at the same time as above using LI-COR 6400XT on intact leaves of whole plants left under the growth-room illumination for 2–4 h after the lights were turned on. Illumination settings in the instrument’s cuvette were set to the same conditions as in the growth room.

### Inhibition of light transduction in detached leaves

Tyrphostin 9 (ENZO, cat. # BML-EI21, 100 mM in DMSO (Sigma, cat # W387520), kept in aliquots at -18 °C) was added to the AXS to a final concentration of 10 μM. AXS with DMSO at a concentration of 100 μl/L (vol./vol.) served as a control in this set of experiments. Excised leaves were kept overnight until the measurements as described above. For surface application, 0.05% Tween 20 (J.T. Baker cat# X251-07) was added to the tyrphostin 9 and control solutions and sprayed prior to overnight perfusion with unmodified AXS. Boxes holding the sprayed leaves did not contain the damp tissue paper, and leaves were patted dry prior to placing them in the light chamber to ensure the leaf surface was dry when the leaves were placed in the LI-COR 6400xt cuvettes.

### Determination of xylem sap pH in detached leaves

#### Leaf preparation for imaging

On the eve of the experiment, leaves from 6-to 7-week-old plants, approximately 2.5 cm long and 1 cm wide, were excised with a sharp blade and perfused (as described) with AXS containing 100 μM of the dual-excitation fluorescent pH probe, FITC-D [fluorescein isothiocyanate conjugated to 10 kD dextran, (Sigma cat. # FD10S), added from a 10 mM stock in double-distilled water]. The excised leaves were then placed in the “double box” setup (i.e., a sealed plastic box inside a dark box) and kept in the dark until measurements the next morning.

#### Dark-treated leaves

Leaves were taken out of the dark boxes and immediately prepared for vein imaging on a microscope slide under low light conditions (<5 μmol m^−2^ sec^−1^).

#### Light-treated leaves

Leaves were taken out of the dark boxes and placed in the growth chamber inside transparent gas sealed boxes (boxes as in K_leaf_) for 30 min of growth room white-light illumination (see above under “Plant growth conditions”) and were subsequently imaged.

The leaves were imaged using an inverted microscope (Olympus-IX8, as detailed by Grunwald et al., 2021). Image capture and image analysis of the intra-xylem pH were as already described (ibid.).

### Determination of the BSCs’ membrane potential (E_M_)

#### Protoplasts preparation

To evaluate the protoplast membrane potential (E_M_), we used a fluorescent (E_M_-sensitive dual-excitation ratiometric dye, Di-8-ANEPPS (Pucihar et al., 2009), and an inverted epifluorescence microscope (Eclipse Ti-S, Nikon, Tokyo, Japan) coupled to an IXON Ultra 888 camera (Andor, UK). A Nikon 40X objective (Plan Fluor 40x/ 0.75, OFN25 DIC M/N2) was used for protoplast viewing and imaging. The experiments were performed between 10 am and 4 pm. BSC protoplasts were isolated as described (Shatil-Cohen et al., 2014) from 6- to 8-week-old SCR:GFP (Col) plants and kept in an Eppendorf vial at room temperature (22 ± 2°C) in the dark until use. A drop with a few protoplasts was added to a bath solution in an experimental chamber, containing 30 μM di-8-ANEPPS and 1% pluronic acid, with or without 10 µM tyrphostin 9 (see “Solutions” below), and protoplasts were allowed to settle for 10 min on a glass coverslip bottom. In separate control assays, we verified the viability of the tyrphostin 9-treated cells by observing that they remained capable of de-esterifying (hydrolyzing) the accumulated SNARF1-AM (see below) and retaining the fluorescent form of the dye, SNARF1, in the cytoplasm.

#### GFP-based selection

An individual, perfectly round BSC (25–32.5 μm diameter) was selected for imaging based on its GFP fluorescence (excitation of 490/15 nm) delivered by a xenon lamp monochromator, Polychrome II (Till Photonics, Munich, Germany), under the control of IW6.1 (Imaging Workbench 6.1) software (Indec Biosystems, Santa Clara, CA), and the emitted fluorescence was filtered via a 515 nm dichroic mirror and a 525/50 nm band-pass barrier filter.

#### Size-based selection

In a single set of experiments, we used a *phot1-5* (Col-gl) mutant and WT (Col-gl) plants that were not labeled with GFP. BS protoplasts are about 50% smaller than mesophyll cells and contain about 50% chloroplast of compared to the mesophyll cells (Shatil-Cohen et al., 2011) and are relatively easy to detect under transmitted light. Thus, we based the BSCs selection on rejecting as “non-BSCs” protoplasts outside the above-mentioned size range (25–32.5 mm diameter). In a separately conducted experiment on protoplasts from GFP-labeled WT (Col), such size-based guesses were confirmed by GFP fluorescence in 68 out of 101 cases.

#### Light treatments and imaging

The selected BSC protoplast in the experimental chamber was exposed to the dark for 10-20 min (control), and a first pair of di-8-ANEPPS fluorescence images was recorded (excitation by a pair of 50 ms pulses, 438 nm then 531 nm, 3 ms apart, was delivered by the Polychrome II; the emitted fluorescence was filtered via a dichroic mirror of 570 nm and 585/20 nm emission band-pass filter; Chroma Technology Corp., Bellows Falls, VT, USA). Subsequently, the protoplast was illuminated for 5-10 min (by a home-made illuminator of crystal clear LEDs from Tal-Mir Electronics, Israel) with either RL (660 nm, 220 µmol m^-2^ s^-1^) or RL + BL (BL: 450 nm, 10–15% of the total intensity of 220 µmol m^-2^ s^-1^) and a second pair of fluorescence images was recorded as above. The images were processed using FIJI (Abràmoff et al., 2004; Schindelin et al., 2012) to yield a pixel-by-pixel fluorescence ratio, as described previously (Wigoda et al., 2017).

#### Fluorescence ratio calibration

BSC protoplasts incubated in di-8-ANEPPS as above were subjected to 5-s-long voltage pulses in the range of +17 to -223 mV using a patch-clamp pipette in a whole-cell configuration. A pair of images was recorded during the second half of each pulse, at a steady state. The patch-clamp imaging setup was as described in detail by Wigoda et al. (2017), except for the substitution of the abovementioned Eclipse microscope and IXON camera for the original image-acquisition instruments. This calibration confirmed the previously published positive correlation between the di-8- ANEPPS fluorescence ratio and depolarization (Supplemental Figure S1). This definitively positive correlation allows us to draw a clear conclusion about the direction of the experimentally induced change in the E_M_, even if the relatively wide data distribution suggests caution in accepting the absolute E_M_ values.

***Solutions*.** <u>Di-8-ANEPPS</u> (Molecular Probes, Life Technologies, cat. # D3167, Oregon, USA): a 10 mM stock solution in DMSO was stored in 10 μL aliquots at -20°C.

<u>Pluronic F-127</u> (Molecular Probes, Life Technologies, cat. # P6867, Oregon, USA), a 20% stock solution in DMSO was stored in 100 uL aliquots at room temperature.

<u>Bath solution (in mM)</u>: 5 KCl, 1 CaCl_2_, 4 MgCl_2_, 10 MES; pH 5.6; osmolality: 435 mOsm, adjusted w/ D-sorbitol.

<u>Patch-clamp pipette solution (in mM)</u>:112 K-gluconate, 28 KCl. 2 MgCl_2_, 10 HEPES, 328

sorbitol, pH 7.5. Osmolarity: 480 mOsm.

<u>Tyrphostin-9</u> was added to the bath solution from aliquots used in the detached-leaves experiments.

### Determination of the BSCs’ cytosolic pH (pH_CYT_)

The experiments were performed between 10 am and 4 pm. BSC protoplasts were isolated according to (Shatil-Cohen et al., 2014) from 6- to 8-week-old SCR:GFP (Col) plants and kept in an Eppendorf vial at room temperature (22 ± 2°C) in the dark until use. The protoplasts were treated as described previously (Torne et al., 2021), with the following modifications: a 10 min treatment was applied, either dark or RL+BL or RL (the illumination setting was as described above for E_M_ determination), the bath solution was used for flushing out the external SNARF-AM during the last 1 min of the dark/light treatments and the cells were imaged within the subsequent 5 min. Image acquisition and analysis to determine the pH_CYT_ values were performed as described previously (Torne et al., 2021).

***Solutions*.** <u>Bath solution</u> for protoplasts’ pH_CYT_ determination: as for E_M_ determination. <u>SNARF1-AM</u>: Carboxy seminaphthorhodafluor-1, acetoxymethyl ester, acetate (Molecular Probes, Thermo Fisher cat. #: C-1272); a 50 µg prepackaged portion was dissolved into a 10 mM stock in DMSO, then stored in 10 μL aliquots at -20°C.

<u>Pluronic F-127:</u> as for E_M_ determination.

### Measuring leaf vein density

Vein density was measured as described previously (Grunwald et al., 2021).

### Determination of the osmotic water permeability coefficient (P_f_) under different light regimes

#### Protoplast isolation and P_f_ determination

These experiments were performed on *SCR:GFP* (Col) protoplasts as described previously (Shatil-Cohen et al., 2014; Grunwald et al., 2021), with minor modifications of equipment and protocol listed in the latter. The P_f_ value was extracted from the image analysis of the rate of the videotaped protoplast challenged with a hypotonic solution, and a simultaneous fit of the swelling time course and the time course of the osmotic concentration change in the bath (C_out_) during the hypotonic challenge. The latter was reconstructed from the videotaped rate of change of optical density of a solution of black ink dissolved in the hypotonic solution flushed into the experimental bath in a separate experiment (Moshelion et al., 2004; Shatil-Cohen et al., 2014; Grunwald et al., 2021).

After their isolation, the protoplasts were divided into two: the “light-treated” protoplasts were kept in a vial under the lights of the growth chamber for the duration of the experiment (between 10 AM and 2 PM), the “dark-treated” protoplasts were kept in a vial in a light-proof box. The two types were sampled alternatingly preserving their respective conditions of illumination (using green safe light while sampling in the dark). The sampled “light-treated“ protoplasts were placed in the experimental chamber and left to settle down for 10-40 min on the bench under a cool-white-led illumination, and then, for 5-7 min on the microscope stage under the unfiltered microscope light (6 V / 30 W halogen lamp, PHILIPS 5761) during preparations and the 1 min videotaping of swelling during the osmotic challenge. The sampled “dark-treated“ protoplasts were placed, under green “safe light”, in the experimental chamber, left to settle for about 10 min and after the brief location of a GFP-positive BSC they remained in total darkness for 10 min. The 1-min videotaping of the protoplast’s swelling during the hypoosmotic challenge in the dark necessitated an illumination by the microscope lamp filtered (to block out the blue light) via a red plastic band-pass filter (∼600-700 nm / 30 μmol m^−2^ sec^−1^).

### Determination of the pH of guttation drops

Whole, intact WT (Col) plants were covered overnight with black plastic bags to increase the relative humidity in their atmosphere and to prevent transpiration. The following morning, immediately after the growth room lights went on, guttation droplets were collected from the tips of the leaves. Ten to twenty guttation droplets were collected and pooled to reach a sample volume of approximately 20 µL in a 200-µL vial, which was immediately sealed. pH was measured within 10 min of sample collection using a micro-pH electrode MI-410 (Microlectrode, Inc., New Hampshire, USA).

### Graphics and Statistics

The Box plots were drawn using ORIGIN (OriginPro, Version 2022. OriginLab Corporation, Northampton, MA, USA). The statistics was performed using JMP (JMP^®^Pro, Version 16.0.0 (512257). SAS Institute Inc., Cary, NC, 1989–2021).

## RESULTS

### phot1 and phot2 blue light receptors are involved in the regulation of K_leaf_

Following the GC model in which the photoreceptors phot1 and phot2 transduce the blue light (BL) stomatal-opening signal, and encouraged by the substantial expression of the BL receptor genes *PHOT1* and *PHOT2* in the transcriptome of Arabidopsis BSCs (Wigoda et al., 2017), we explored the roles of these two light receptors in the regulation of leaf hydraulic conductance (K_leaf_). We compared the K_leaf_ of WT plants to the K_leaf_ of knockout mutants lacking one or both light receptors (*phot1-5*, *phot 2-1* and *phot1-5 phot 2-1*; Figure 1) under two light regimes, red light (RL) and RL+BL. Under RL+BL, the K_leaf_ of WT leaves was more than 3-fold higher than it was under RL treatment and more than 2-fold higher than the K_leaf_ values of the three mutants treated with RL+BL. Also, under RL+BL, the K_leaf_ of the *phot2-1* mutant was about 40% higher than it was under the RL treatment. In contrast, BL did not seem to increase the K_leaf_ of the *phot1-5* mutant or that of the double mutant. Under RL, there were no differences between the K_leaf_ levels of the different mutants. However, while the WT’s K_leaf_ was no different from that of the single-mutants, it was about double that of the double mutant (Figure 1). The BL response was particularly depressed in the double mutant: only 13% of the WT’s K_leaf_ BL response remained in the double mutant, as compared to 32% and 40% of the WT’s K_leaf_ in *phot1*-*5* and in *phot2*-*1*, respectively (Figure 1), and only 35% of the WT’s g_S_ BL response remained in the double mutant, as compared to 57% and 71% of the WT’s g_S_ in *phot1*-*5* and *phot2*-*1*, respectively (Supplemental Figure S2A).

**Figure 1.**
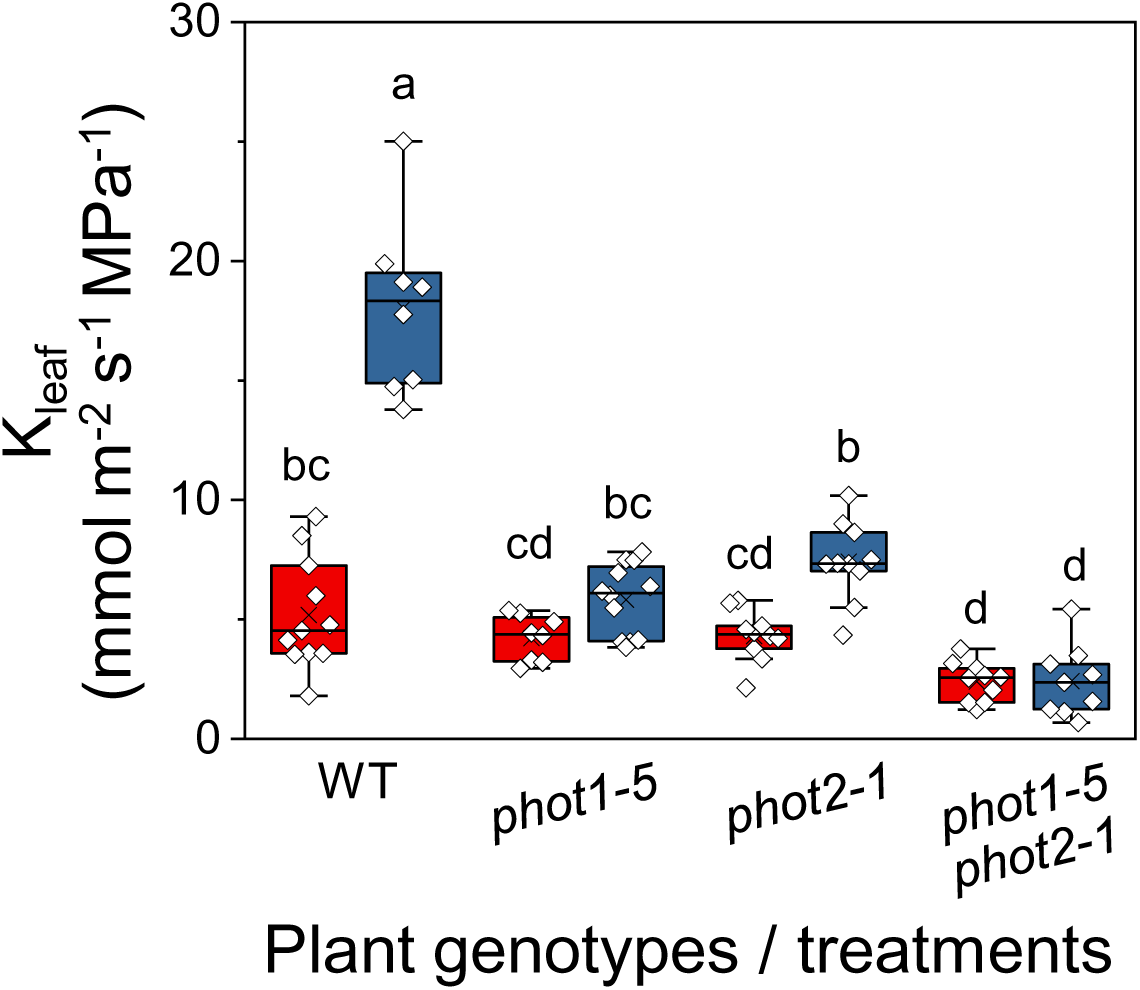
The effect of blue light on the K_leaf_ of PHOT receptor mutants. K_leaf_ of fully expanded, excised leaves of WT (Col-gl) and *PHOT* mutants [*phot1-5* (Col-gl), *phot2-1* (Col-gl) and a double mutant, *phot1-5 phot2-1* (Col-gl)] determined immediately after 15 min of illumination with red light (RL, red box; 220 µmol m^−2^ s^−1^) or red + blue light (RL+BL, blue box; 220 µmol m^−2^ s^−1^, consisting of 90% RL and 10% BL). The box height delimits 25-75 % of the data range, the symbols – the individual data values, the solid line – the median, x – the mean, and the whiskers delimit ±1.5 times the inter-quartile range (IQR). Different letters indicate statistically different means (ANOVA, Tukey-HSD test, P<0.05). Note the reduced response to RL+BL in the mutant lines.

The higher calculated K_leaf_ value of the WT under BL (as compared to RL) was due to a greatly increased (less negative) leaf water potential, Ψ_leaf_, and an appreciably higher transpiration rate, E (Eq. 1, Materials and methods). This invites the interpretation that the BL induced a highly conductive water pathway into the leaf, enabling higher radial water influx which, in turn, was able to compensate for the BL’s enhancement of E (i.e., water efflux from the leaf via the BL-increased g_S_; Supplemental Figure S2).

### BSC-specific silencing of phot1 and phot2 receptors decreased K_leaf_

To examine the specific participation of the BSCs’ light receptors in the BL-induced increase in K_leaf_, we silenced either the *PHOT1* or the *PHOT2* gene using amiRNA (artificial microRNA) under the BSC-specific promoter, *SCARECROW* (*SCR*, see Materials and Methods). Under RL+BL illumination, the K_leaf_ values of leaves of both types of transgenic plants, *SCR:amiR-PHOT1* and *SCR:amiR-PHOT2*, were lower than the K_leaf_ values of the WT leaves (by 40–50%; Figure 2). In contrast, all of the *SCR:amiR-PHOT* plants had WT-like g_S_ and E, while their Ψ_leaf_ levels were lower than that of the WT (by 50–90%), suggesting a reduction in the radial influx of water into the leaf (Supplemental Figure S3).

**Figure 2.**
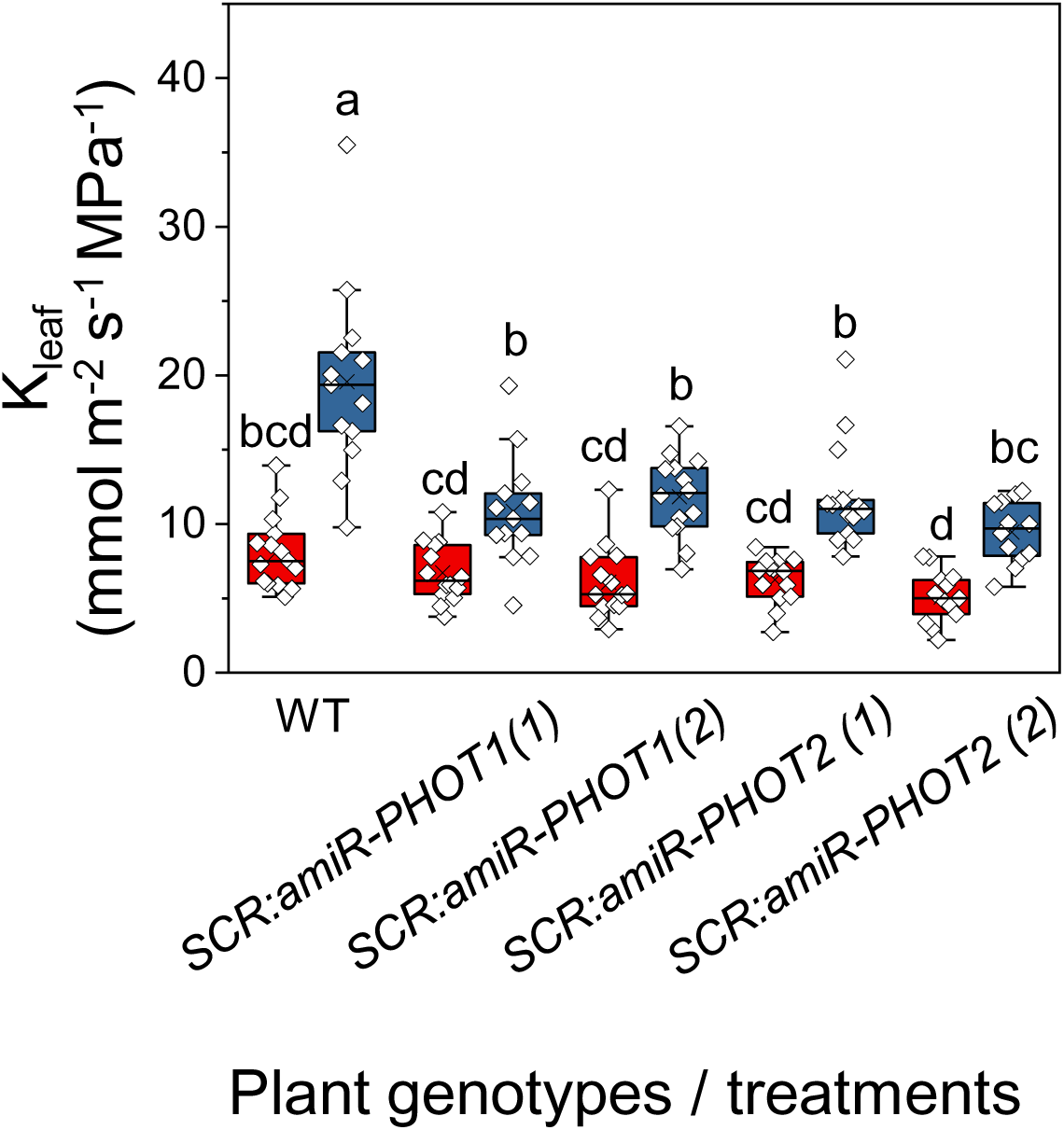
*amiR*-silencing of *PHOT1* or *PHOT2* (*SCR:amiR-PHOT* plants) in the BSCs reduces K_leaf_ under RL+BL light. K_leaf_ was determined in fully expanded leaves of WT, *SCR:amiR-PHOT1* (two independent lines 1,2) and *SCR:amiR-PHOT2* (two independent lines 1,2) plants treated with RL or RL+BL (as in Figure 1). Other details are also as in Figure 1.

### BSC-specific complementation of the *phot1-5* mutant with *PHOT1* restored K_leaf_

We complemented the *phot1-5* (Col-gl) mutant plants with *SCR*:*PHOT1* to restore phot1 activity exclusively in the BSCs (Materials and methods) and compared them to WT (Col-gl) plants and to *phot1-5* plants. *SCR*:*PHOT1* plants illuminated with RL+BL had their K_leaf_ restored from *phot1-5* values to WT levels (Figure 3). While, as expected, BL increased the WT’s g_S_, the mutant’s g_S_ and the g_S_ of the complemented mutant did not change under BL, also as expected from the main BL-receptor-devoid guard cells (Supplemental Figure S4). Under BL, the Ψ_leaf_ of the *SCR*:*PHOT1*-complemented mutant was restored to the WT level and both of those values were more than 50% greater than the mutant’s Ψ_leaf_ (Supplemental Figure S4). Thus restored Ψ_leaf_ was the major contribution to the restored high K_leaf_ of the complemented mutant, suggesting a restoration of radial water influx to the leaf.

**Figure 3.**
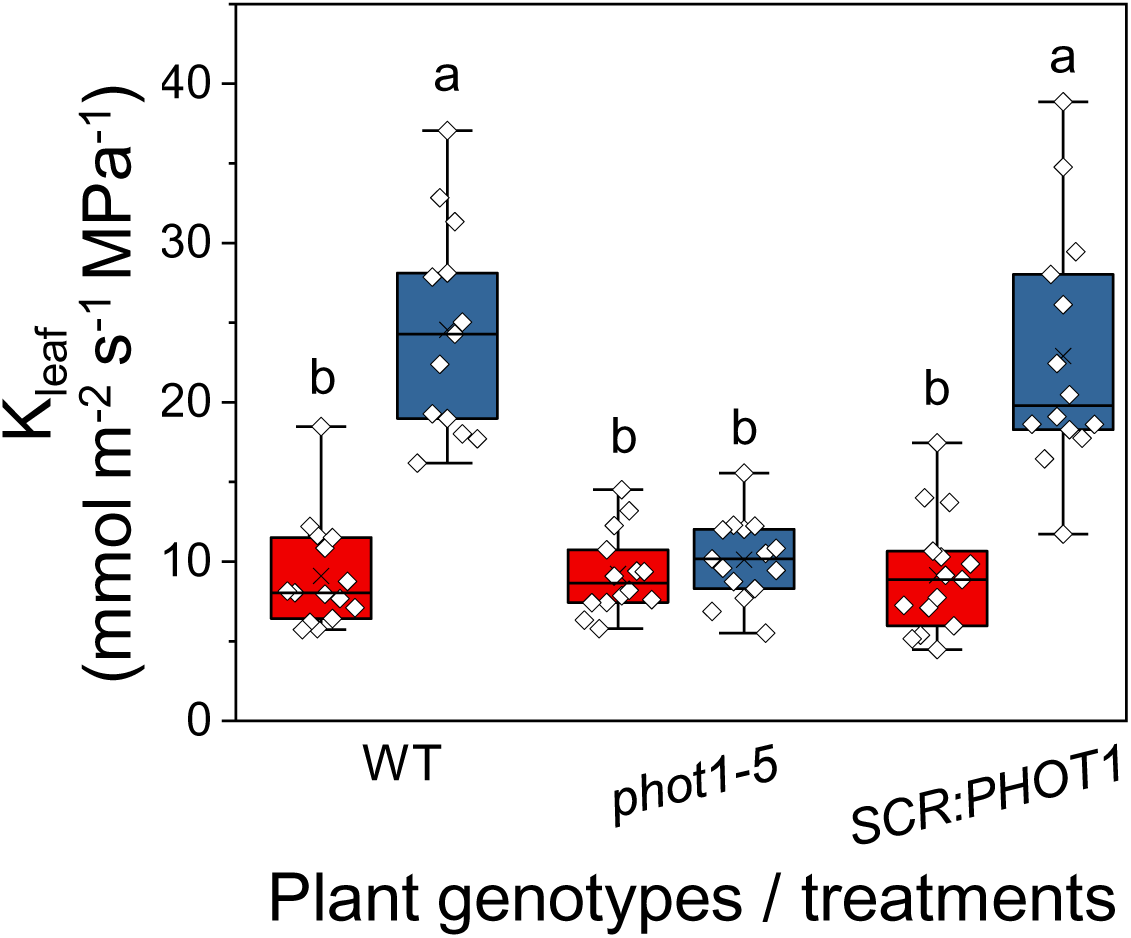
BSC-directed *PHOT1* complementation of the *phot1-5* mutant restores the normal K_leaf_. Fully expanded leaves of WT (Col-gl), *phot1-5* (Col-gl) and *phot1-5* (Col-gl) plants complemented with *SCR:PHOT1* were subjected to illumination treatments, as in Figure 1. Other details are also as in Figure 1.

### The kinase inhibitor, tyrphostin 9, prevented the increase in K_leaf_ under blue light

The kinase inhibitor tyrphostin 9 suppresses BL-dependent H^+^-ATPase phosphorylation (of the penultimate threonine) in guard cells (Hayashi et al., 2017). To examine further whether the vascular phot receptors initiate a similarly sensitive phosphorylating event in the BL-dependent K_leaf_ regulation pathway, we perfused detached WT leaves with AXS, with or without 10 μM tyrphostin 9, and exposed them to RL or RL+ BL. The K_leaf_ values of RL+BL-illuminated leaves perfused with tyrphostin 9 were about 50% lower than those of RL+BL -illuminated control leaves (without tyrphostin 9), and were no different than the K_leaf_ of leaves illuminated with only RL (with or without exposure to tyrphostin 9; Figure 4A, left panel, Supplemental Figure S5). However, g_S_ was not affected by the petiole-delivered tyrphostin 9 (Figure 4B, left panel). In contrast to the perfused tyrphostin 9, spraying the inhibitor on the leaf surface did not interfere with the almost 3-fold increase in K_leaf_ that was induced by BL (Fig. 4A, right panel), but it did prevent the BL-induced increase in g_S_ (Figure 4B, right panel). The impaired stomatal response as a result of the application of tyrphostin 9 directly to the leaf surface is in agreement with tyrphostin 9’s previously reported inhibition of BL-induced stomatal opening in epidermal peels (Hayashi et al., 2017). Leaf transpiration, E, behaved in a pattern similar to g_S_ (Supplemental Figure S6A). Petiole-fed tyrphostin 9 abolished the BL-induced increase in Ψ_leaf_, but tyrphostin 9 sprayed on the leaf surface did not prevent it (Supplemental Figure S6B).

**Figure 4.**
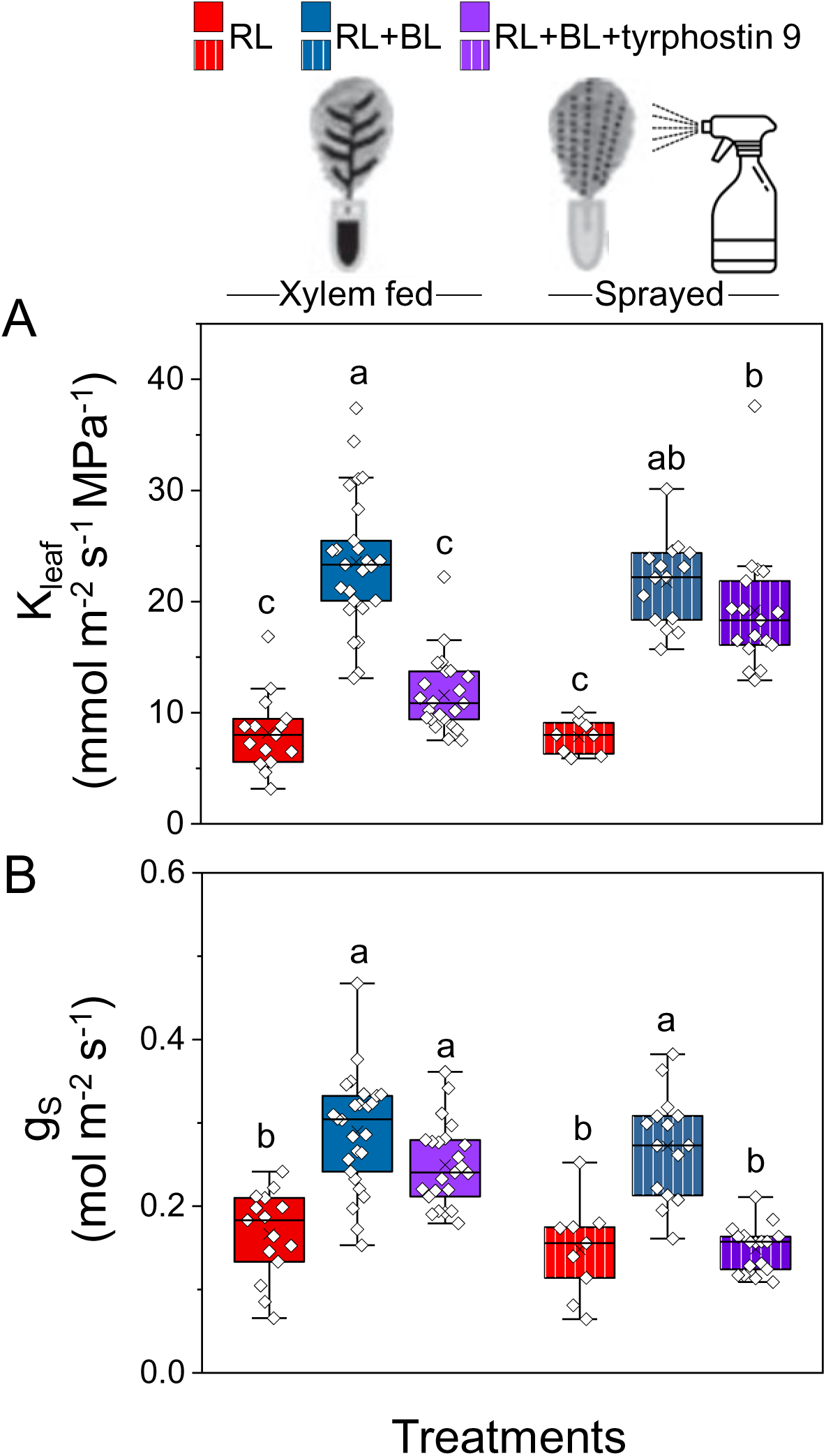
The kinase inhibitor tyrphostin 9 either abolished the BL-induced K_leaf_ increase or prevented the BL-induced stomatal opening - depending on the treatment route. Fully expanded leaves of WT (Col-gl) were pre-incubated (petiole deep) in AXS without or with tyrphostin 9 (10 μM) or sprayed with AXS with or without tyrphostin 9 (10 μM) and kept overnight in dark boxes. Immediately after dark, leaves were illuminated for 15 min with RL or RL+BL, as indicated. (A) K_leaf_. (B) Stomatal conductance (g_S_). Other details are as in Fig. 1. Note that K_leaf_ was reduced when tyrphostin 9 was fed via the petiole, but not when it was sprayed on the plants, and that the opposite was true for g_S._

### Blue light hyperpolarizes the BSCs and raises their cytosolic pH

To investigate further whether the BSCs’ BL signal transduction pathway stimulates their plasma-membrane H^+^-ATPases (as in GCs), we monitored the membrane potential of BSC protoplasts using the potentiometric dual-excitation dye di-8-ANEPPS. Indeed, 5–10 min of RL+BL illumination hyperpolarized the BSC protoplasts relative to RL alone or to the dark (absence of illumination), while 10 µM tyrphostin 9 added in the bath depolarized the BSCs irrespectively of RL+BL illumination or its absence (Figure 5A, Supplemental Figure S1), hence, apparently independently of the inhibition of AHA activity.

**Figure 5.**
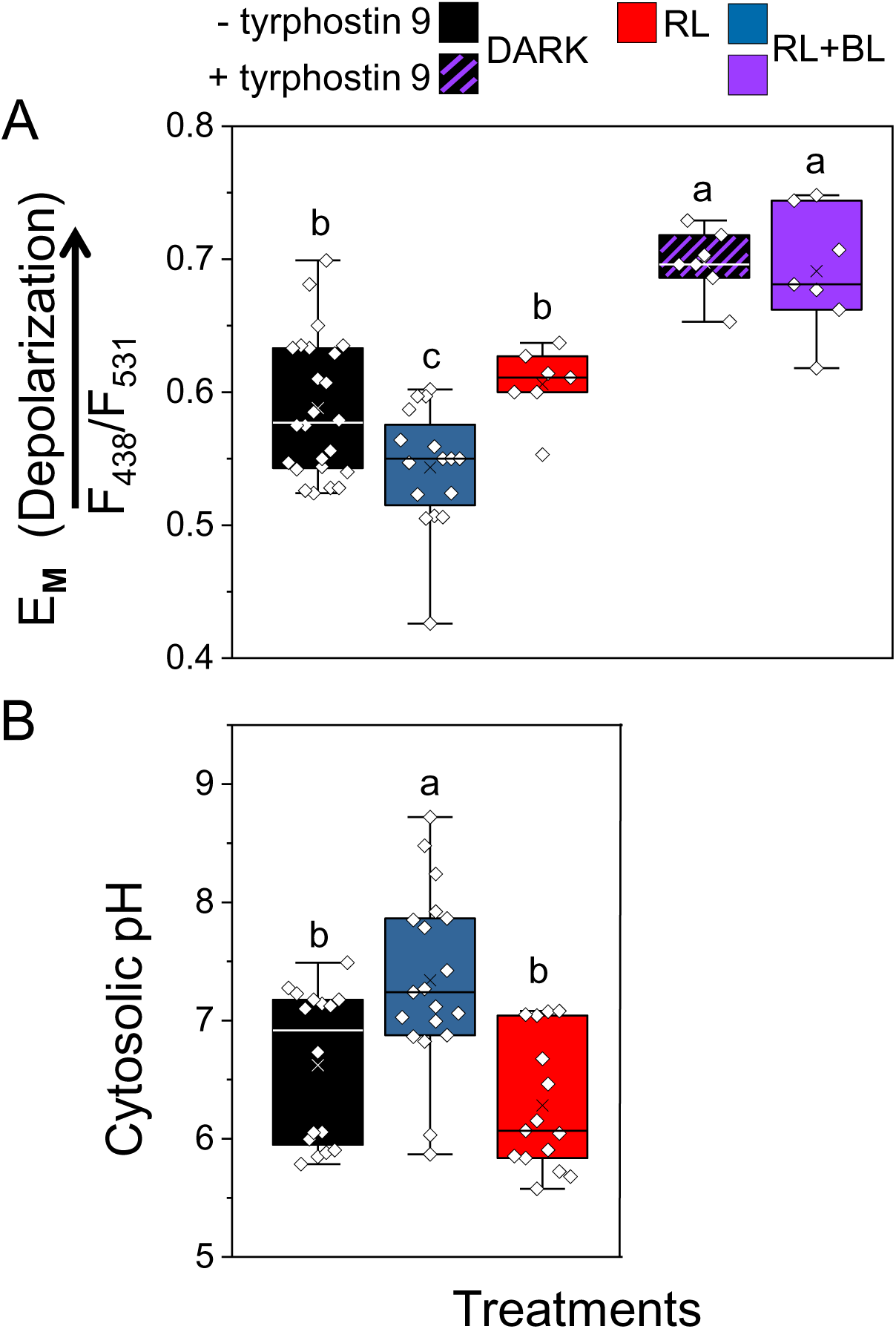
Blue light hyperpolarizes the isolated BSC protoplasts and alkalinizes their cytosol relative to red light or darkness, and tyrphostin 9 depolarizes them. (A) Membrane potential (E_M_, in units of the fluorescence ratio) of BSC protoplasts, with or without bath-added tyrphostin 9 (10 µM), determined using the dual-excitation, ratiometric fluorescent dye di-8-ANEPPS. After at least 10 min in the dark, the cells were immediately imaged (dark treatment) or illuminated for an additional 5–10 min either with RL or RL+BL and then imaged (Materials and Methods). The symbols represent the mean fluorescence ratio (F_438_/F_531_, F-ratio) values from individual protoplasts (biological repeats), derived from analysis of their images captured at the indicated excitation wavelengths in three to eight independent experiments. The positive correlation between E_M_ and the F-ratio (with depolarization indicated by the arrow) was verified in separate experiments using patch clamp (Supplemental Figure S1). Different letters denote significantly different values (ANOVA, post hoc Tukey’s test; P < 0.05). Other box plot details are as in Fig. 1 (B) Cytosolic pH of BSC protoplasts determined using the fluorescent dual-emission pH probe SNARF1. Protoplasts were dark-treated for 10 min and then further treated with 10 min of either dark, RL or RL+BL, then imaged. Cytosolic pH values were derived from the analysis of the captured pairs of images of individual protoplasts (biological repeats); the emitted fluorescence ratio (F640/F585) values were converted to pH values based on a calibration curve (as detailed in Torne et al., 2021). The data are from at least three independent experiments. Other details are as in A.

To further explore our hypothesis of H^+^-ATPase activation by BL, we monitored the cytosolic pH (pH_CYT_) using the ratiometric, dual-emission fluorescent pH-sensitive dye, SNARF1. The pH_CYT_ of BSCs illuminated for 10 min with RL+BL (following dark treatment) was significantly more alkaline (by 0.72 pH units) than the pH_CYT_ of dark-treated BSCs and 1.06 pH units higher than the pH_CYT_ of the BSCs that were treated with only RL (Figure 5B), consistent with BL-induced activity of H^+^-ATPases, which move protons from the BSCs into the xylem.

### AHA2 plays a role in the blue light-induced regulation of K_leaf_

Recently, we reported that AHA2 plays a major role in K_leaf_ regulation by acidifying the xylem sap. To further examine whether the BSCs’ H^+^-ATPase AHA2 participates in the *BL-initiated* K_leaf_ regulation, we examined the AHA2 knockout mutant *aha2-4* and its BSC-complemented transgenic plant line *SCR*:*AHA2* [i.e., the *aha2-4* mutant with AHA2 expressed only in its BSCs (Grunwald et al., 2021)] under the two light regimes (RL and RL+BL). The K_leaf_ of *aha2-4* leaves did not respond to the RL+BL illumination and was only about 40% of the K_leaf_ of BL-treated WT leaves. However, similar illumination increased the K_leaf_ of the AHA2-complemented (*SCR*:*AHA2*) plants to almost 65% of the WT’s K_leaf_ (Figure 6), indicating that the BL-induced increase in K_leaf_ is dependent on AHA2. In contrast, under RL+BL, the E and g_S_ of *SCR*:*AHA2* did not differ from the E and g_S_ of the *aha2-4* mutant, and they remained unaffected by BL and lower than the E and g_S_ of the WT (Supplemental Figure S7A,B). Notably, the Ψ_leaf_ of *SCR*:*AHA2* was about 40% higher than the Ψ_leaf_ of *aha2-4* and did not differ from that of the WT, suggesting a restoration of radial water influx to the leaf (Supplemental Figure S7C).

**Figure 6.**
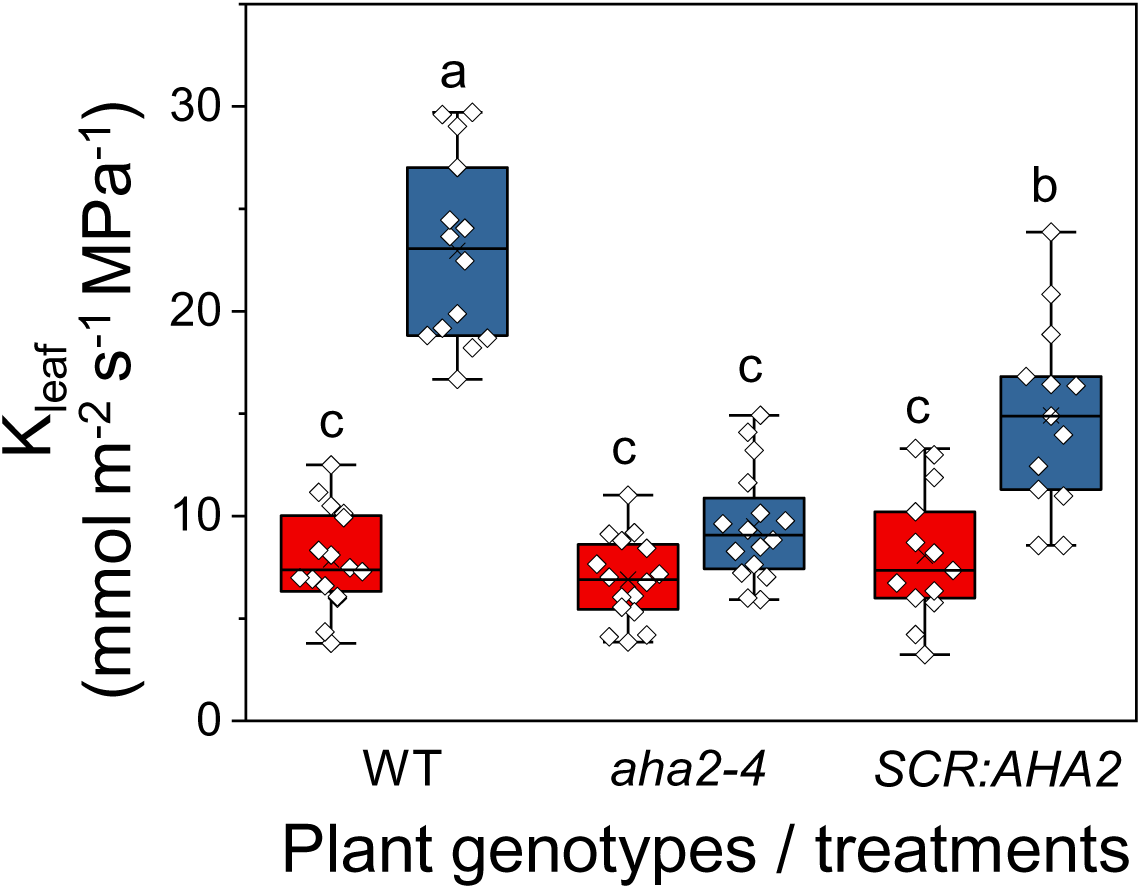
The BSC H^+^-ATPase, AHA2, mediates the BL-induced increase in K_leaf_. Fully expanded excised leaves of WT (Col0) plants, *aha2-4* mutant plants (Col0) and *aha2-4* plants complemented in their BSCs with *SCR:AHA2* were illuminated for 15 min immediately after dark, as in Figure 1. The box-plot details are also as in Figure 1. Note the marked restoration of K_leaf_ in the *aha2* mutant plants complemented solely in their BSCs.

### Morning light after night darkness acidifies the xylem sap

To delve further into the acidification-mediated link between light and K_leaf_, we explored the diurnal transition between the xylem sap pH in the night and in the morning. We hypothesized that the xylem sap pH would be reflected in the pH of guttation drops exuded via hydathodes from the ends of the xylem conduits. Therefore, to take a preliminary peek at the xylem sap pH at night, we measured the pH of guttation drops collected from the tips of WT leaves before the lights were turned on and found that their mean pH was 7.2 ± 0.04 (±SE, 15 biological repeats collected over 3 days). Next, we measured the pH of the xylem sap in detached leaves, comparing leaves before dawn to leaves after dawn (i.e., leaves at the end of an overnight dark treatment to leaves after a 30 min of illumination with white light in the growth chamber). This comparison included leaves of WT, *phot1-5* and *SCR:PHOT1* plants. Illumination acidified the xylem sap in WT plants by approx. 0.6 pH units (from 6.0 to 5.4) and in the *SCR: PHOT1* plants by approx. 0.8 pH units (from 6.1 to 5.3), while the xylem pH in *phot1-5* remained unchanged, at about 5.9 (Figure 7), suggesting that phot1 in the BSCs is necessary for light-activated xylem acidification, which, in turn, increases K_leaf_ (Grunwald et al., 2021).

**Figure 7.**
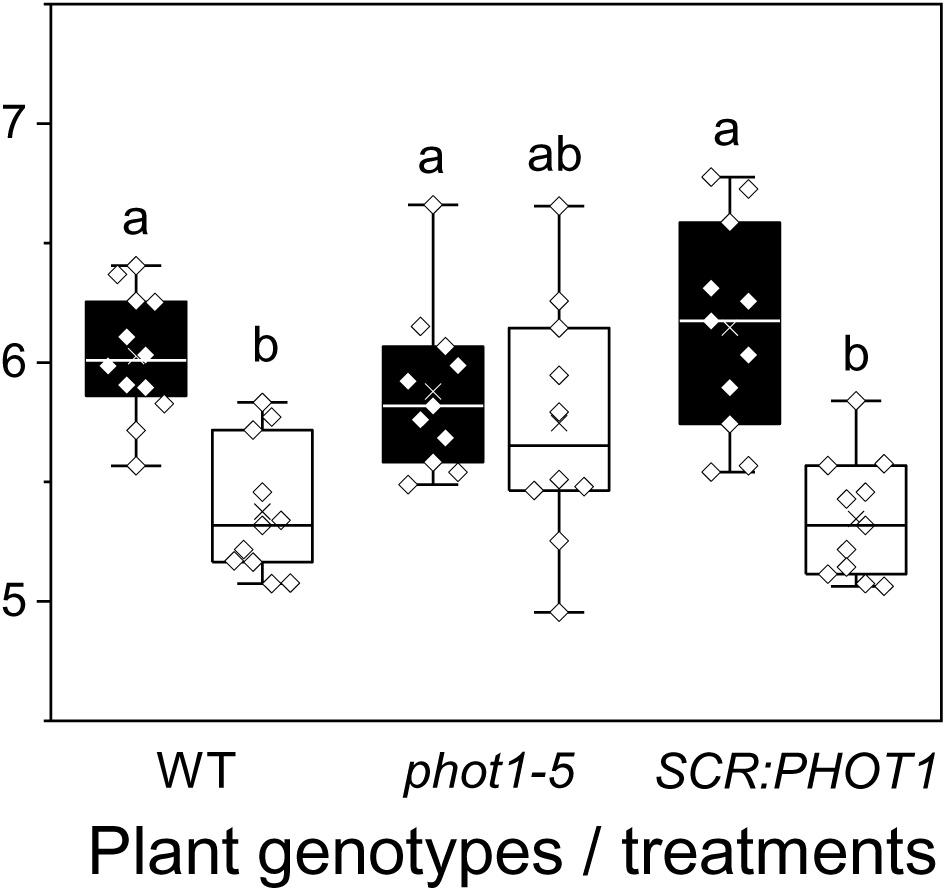
Illumination with white light for 30 min acidified the xylem pH of WT plants and *SCR*:*PHOT1*-complemented mutants, but not that of the *phot1-5* mutant. The leaves of WT (Col-gl) plants, *phot1-5* (Col-gl) mutants and the *SCR:PHOT1*-complemented *phot1-5* mutant (as in Figure 3) were illuminated with white light in the growth room (150–200 μmol m^−2^ sec^−1^). Black boxes indicate dark, white – white light. Other box plot details are as in Fig. 1.

### Light increases the osmotic water permeability, P_f_, of the BSCs

To explore the link between light and K_leaf_ at the cellular level, we tested whether light affects the osmotic water permeability (P_f_) of the BSC membrane. We determined the P_f_ of isolated BSC protoplasts from videotapes of their swelling during a hypotonic challenge after illumination with white light or after 10 min in the dark. The illuminated BSCs had higher P_f_ than the dark-treated ones (Figure 8), suggesting that the morning increase in the movement of water from the xylem to the BSCs (and then, presumably, further into the mesophyll) is due to increased permeability of the BSC membranes to water.

**Figure 8.**
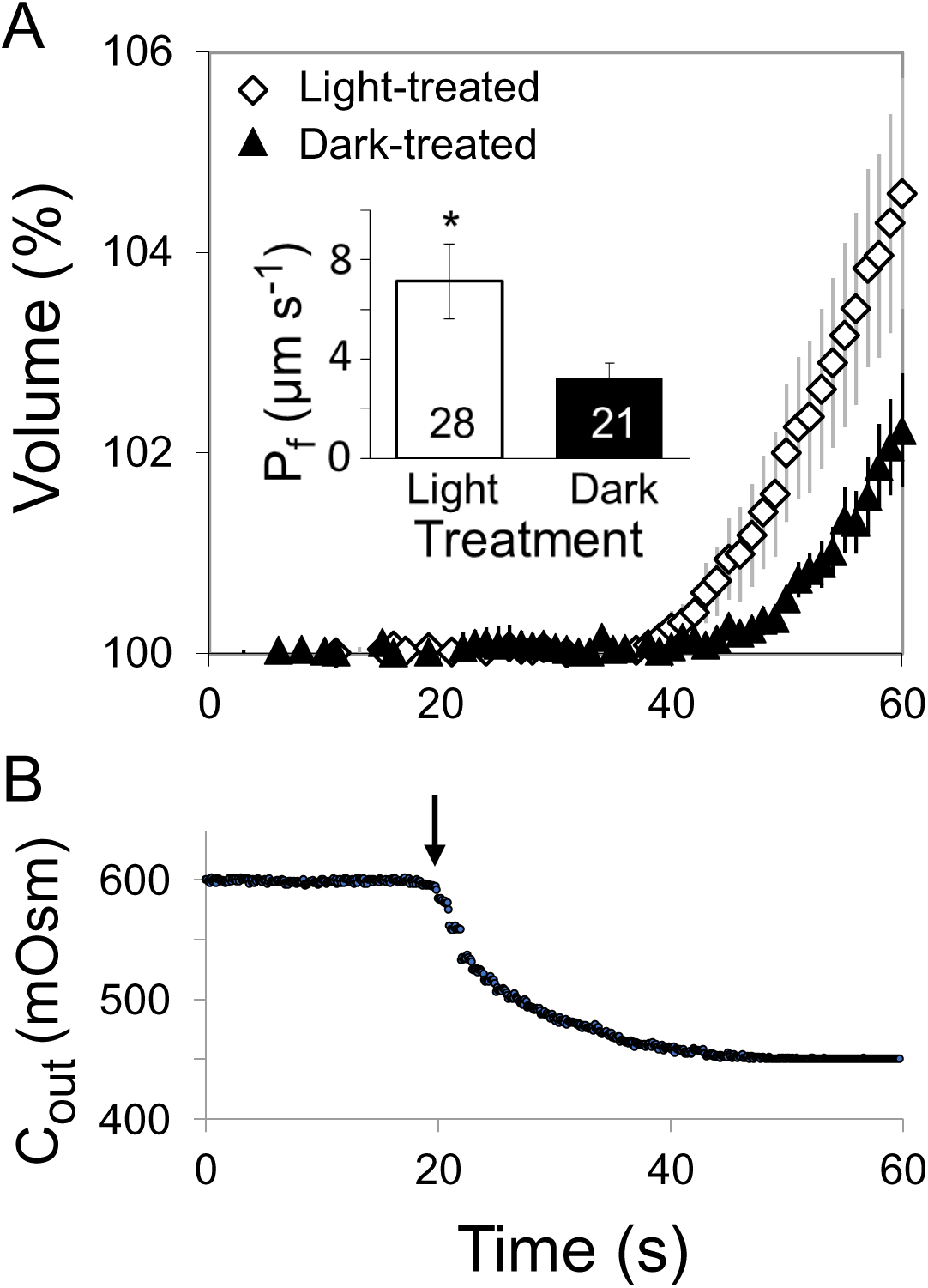
White light increases the BSCs’ membrane osmotic water permeability coefficient (P_f_). (A) Time course of the swelling (mean relative volume ±SE vs. time) of n bundle sheath protoplasts from *SCR:GFP* plants upon exposure to a hypotonic solution under the same illumination conditions (white light or darkness) as during the light treatment preceding the assay (Materials and Methods). Inset: the mean (±SE) initial P_f_ values of the indicated number (*n)* of BSC protoplasts under two illumination regimes from five independent experiments. The asterisk indicates a significant difference (at *P* < 0.05; Student’s two-tailed, non-paired *t*-test). (B) Time course of the osmotic concentration change in the bath (C_out_) during the hypotonic challenge, the arrow indicates the onset of the bath flush (Materials and Methods).

### Leaf vein density is not affected by the expression of phot1 or phot2

To resolve whether the diminished K_leaf_ of the phot receptor mutants that we observed under BL was due to differences in leaf vein density, we quantified the vein density of all of the genotypes that we studied and found no differences among them (Supplemental Figure S8).

## DISCUSSION

### Blue-light signal transduction in BSCs is reflected in the K_leaf_ increase

We demonstrate here an autonomous BL signaling mechanism in Arabidopsis BSCs that underlies the light-induced increase of K_leaf_, resembling the “classical” short-term, BL-induced signaling pathway for stomatal opening mediated by the photoreceptors phot1 and phot2 (Zeiger et al., 1983; Sakai et al., 2001; Kinoshita et al., 2001). To explain the underlying mechanism, we built on our earlier findings that the receptor genes *PHOT1* and *PHOT2* are substantially expressed in BSCs (Wigoda et al., 2017; Supplemental Table 2). Here, we link these findings using the phot receptor mutants *phot1-5* and *phot2-1* and the double mutant *phot1-5 phot2-1*. Based on the complete abolishment of the BL-K_leaf_ response (relative to RL alone) in the *phot1*-*5* mutant, which is presumed to possess phot2 activity, phot2 would appear completely redundant, were it not for (a) the strong depression of the BL-K_leaf_ response – compared to WT response – in the *phot2-1* mutant, presumed to possess phot1 activity, and (b) even more severe impairment of the double mutant’s BL-K_leaf_ response compared to the responses of both single mutants (Figure 1).

Also the BL-g_S_ responses that we observed in the three phot mutants were depressed to different relative degrees resembling the gradation of BL-K_leaf_ responses (Supplemental Figure S2A) and suggesting a similar dependence of BL signaling on phots in the two systems (BSCs and GCs). Interestingly, after 2-4 h of illumination (well beyond our usual 20 min of illumination) the *phot* single mutants’s g_S_ re-acquired the WT-like sensitivity to BL (Supplemental Figure S9), similar to an earlier report on stomatal aperture after 2-4 h of illumination with BL (Kinoshita et al., 2001). This could perhaps reflect a gradually accumulating effect of the GCs’ AHA activity, overcoming the partially impaired phot-to-AHA signal transduction in the single *phot* mutants.

That the phots (and particularly phot1) are indispensable to BL signaling in the BSCs, as they are in GCs (Kinoshita et al., 2001), is additionally emphasized by our two BSCs-delimited genetic phot manipulations: (a) the severe impairment of the BL-induced K_leaf_ increase in WT plants in which either phot1 or phot2 was silenced by the respective *SCR:amiR-PHOT* (Figure 2) and (b) the full restoration of K_leaf_ sensitivity to BL by *phot1-5* complementation with *SCR:PHOT1* (Figure 3). These manipulations, which did not affect g_S_ (Supplemental Figures S3A, S4A), also emphasize the independence of the BSCs’ BL signal transduction from that of the GCs.

Prompted by finding BLUS1, the phots’ indispensable partner kinase in the BL-induced early-signaling complex of GCs (Takemiya et al., 2013; Hayashi et al., 2017) also in the BSCs transcriptome (Wigoda et al., 2017; Supplemental Table S2), we found that the *blus1* mutant’s K_leaf_ does not respond to BL (Supplemental Figure S10), underscoring further the resemblance in BL signaling between the two systems.

Additionally, the tissue-localized effects of the kinase inhibitor, tyrphostin 9, which presumably inhibited a BL-signaling kinase downstream of phot-BLUS1, demonstrated both the similarity between the BSCs’ and the GCs’ BL signaling pathways in WT plants, and, again, the independence of the two systems. Both signaling pathways were inhibited by tyrphostin 9, but only when it was applied in close proximity to the target tissue: The BSC system was affected only by the petiole-fed tyrphostin 9, which inhibited the BL-induced increase of K_leaf_ and Ψ_leaf_ without affecting g_S_; whereas the GC system was affected only when tyrphostin 9 was sprayed on the leaf surface (i.e., on GCs), which abolished the BL-induced g_S_ increase without affecting K_leaf_ (Figure 4, Supplemental Figure S6). Notwithstanding the above, the depolarization of BSCs by tyrphostin 9 both under RL+BL and in the dark (Fig. 5A), suggests an additional or an alternative underlying mechanism *outside* the AHA-activating BL-signaling pathway.

Recently, we reported that the BSC H^+^-ATPase AHA2 plays a major role in the acidification of the xylem sap, which, in turn, increases K_leaf_ (Grunwald et al., 2021). Indeed, the H^+^-ATPase mutant, *aha2-4* had a diminished (compared to WT) BL-induced increase in K_leaf_ (Figure 6; likely due to a more alkaline xylem pH; Grunwald et al., 2021), as well as a diminished BL-induced increase in g_S_ (Supplemental Figure S7). In contrast, *aha2-4* plants complemented with the *SCR:AHA2* – even though directed only to BSCs – had their BL-induced K_leaf_ increase restored (Figure 6), while their g_S_ remained unaffected (Supplemental Figure S7), affirming the importance of the BSC AHA2 in mediating the BL-induced increase in K_leaf_ independently of g_S_ (i.e., of GCs).

Thus, with the genetic manipulation of the BL receptors phot1 and phot2 in the BSCs, the tissue-differential responses to pharmacological treatment and genetic manipulation of the AHA2 we demonstrated the independence of the BSCs’ BL signaling from the GCs’ BL signaling. This notion of independence is particularly important since the BSC layer constitutes a hydraulic valve between the xylem and the leaf, regulating water loss to the atmosphere *in series* with stomatal apertures.

### Blue-light signal transduction in BSCs is reflected in their H^+^-ATPase activation

Four of our findings focus on the activation of H^+^-ATPase as the end-point of BL signaling: at the level of single isolated BSCs, (a) the BL-induced alkalinization of the BSCs cytosol (relative to dark or to RL; Figure 5B), (b) the BL-induced hyperpolarization in WT (Col) BSC protoplasts (relative to dark or to RL, Figure 5A), and also in WT (Col-gl) BSC protoplasts (Supplemental Figure S11A, Materials and Methods), (c) the absence of hyperpolarization under RL+BL treatment in the BSCs protoplasts of the *phot1-5* mutant and its restoration in the *phot1-5* mutant complemented with *PHOT1* solely in its BSCs (Supplemental Figure S11B); at the level of the detached leaf, (d) acidification of the xylem sap by white light (relative to dark) in WT, the absence of the white light response in the *phot1-5* mutant and a restoration of this response in the abovementioned *PHOT1-*complemented mutant (Figure 7). Taken together, both types of the pH responses (which occur by proton extrusion from the cytosol) and the hyperpolarization (which could also result from proton extrusion via the plasma membrane, resembling the hyperpolarization of GCs in response to BL (Assmann et al., 1985; Roelfsema et al., 2001), are strong evidence linking the phot receptors with the BL-induced activation of the BSCs’ AHA2 in a similar way to BL activation of the GCs’ AHA (Hayashi et al., 2017).

### Light induces the increase of BSCs’ membrane water permeability, K_leaf_ and K_ros_

Here, we report that in addition to external (experimentally imposed and constant) low pH, light also increases the P_f_ of isolated BSCs (Figure 8). Evidence is accumulating that P_f_ level reflects the underlying activity of aquaporins. For example, frog oocytes expressing AtPIP2;1 had higher P_f_ induced by co-expressing two of the Arabidopsis 14-3-3 protein isoforms which bound to the AtPIP2;1 (Prado et al., 2019). Thus, it would be tempting to speculate that light increased the BSCs’ P_f_ (Figure 8) by enhancing the activity of one or more of the 11 BSCs’ aquaporins (Supplemental Table S2).

Alternatively, or in addition, the light-induced increase in BSC P_f_ could be due to the alkalization of the BSCs’ cytosol (Figure 5B), sensed by a histidine (His 197, conserved in all Arabidopsis PIP aquaporins; assayed in AtPIP2;2; Tournaire-Roux et al., 2003) and a leucine (Leu206 in BvPIP2;2 Fortuna et al., 2019) in the cytosol-facing D loop of the aquaporin protein. Moreover, the concurrent hyperpolarization of BSCs by BL (Figure 5A) could perhaps also contribute to the activation of the BSCs’ aquaporins, as suggested by *in silico* simulations (Hub et al., 2010).

The role of aquaporins in controlling K_leaf_ was identified previously [in walnut, Ben Baaziz et al., 2012; in grapevine (*Vitis vinifera*), Vitali et al., 2016; in poplar (*Populus trichocarpa*), Muries et al., 2019; in Arabidopsis BSCs, Sade et al., 2014; (Prado et al., 2013)]. In particular, Prado et al. (2019) demonstrated a contribution of the abundant – also in the leaf veins – aquaporin AtPIP2;1 to the diurnal and circadian rhythms of the hydraulic conductance of the Arabidopsis rosette, K_ros_, which peaked before midday. Thus, in addition to the likely P_f_-promoting effect of the slightly acidic pH (6) perfusion solution (a

link established by Grunwald et al., 2021), that Prado et al., (2019) used for their K_ros_ measurements, the changes that they observed in K_ros_ also reflected the effects of light and circadian rhythms (Prado et al., 2019). In particular, the circadian regulation of the AtPIP2;1 by its protein-protein interaction with a 14-3-3 protein (Prado et al., 2019), which resembles the multitude of protein-protein interactions regulating mammalian aquaporins (reviewed by Roche and Törnroth-Horsefield, 2017), suggests an additional possibility of BL regulation of the BSCs’ aquaporins: directly via a protein-protein interaction with the phots.

### BL signaling increases the K_leaf_ -a model

Whatever the exact underlying molecular mechanism, our results support the following model of K_leaf_ increase by BL, in a GC-like, BL signal transduction pathway: the BSCs’ phot1 and phot2 perceive the BL, which, via BLUS1 and unknown downstream protein kinases (the first of which could perhaps be BHP, (Hayashi et al., 2017; Hosotani et al., 2021) activates the plasma membrane H^+^-ATPase (AHA2), resulting in the hyperpolarization of the BSC, the alkalization of the BCSs’ cytosol and the acidification of the xylem sap, leading to an increase in the BSCs’ P_f_ and a subsequent increase in K_leaf_.

Expressed schematically:

BL → activation of BSC *phot*s → phosphorylation* & activation of BLUS1 and downstream kinases → activation of BSC H^+^-ATPase (AHA2) → hyperpolarization and cytosol alkalization of BSCs and acidification of xylem sap → increase of P_f_ of BSCs → K_leaf_ increase

*We hypothesize that the abovementioned phosphorylation steps are similar to those seen in GCs (Figure 9), but this has yet to be explored in BSCs.

**Figure 9.**
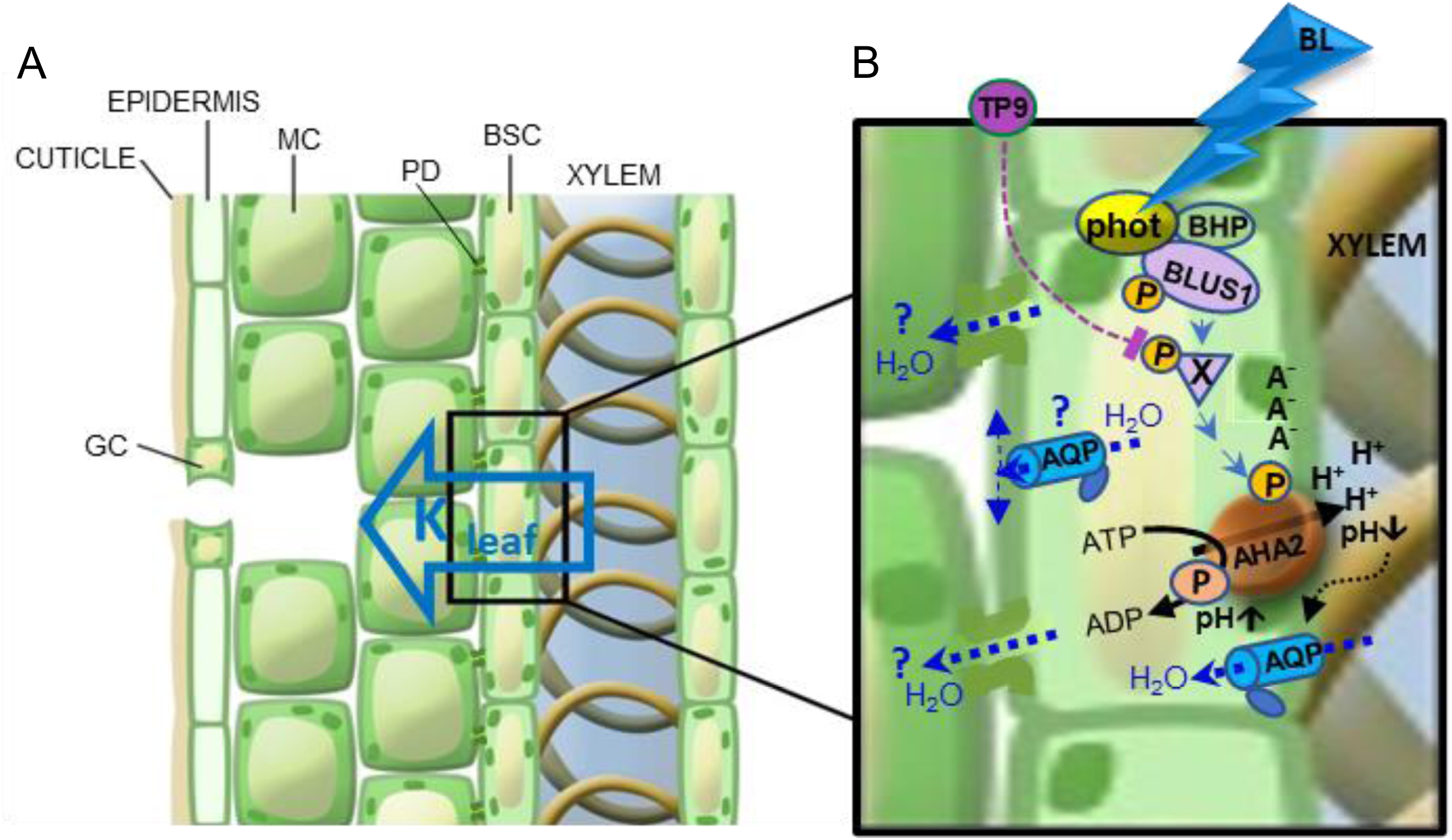
Proposed BSC-autonomous blue-light signal transduction pathway regulating K_leaf_, the leaf hydraulic conductance – an artist’s rendering. (A) A leaf radial water path, from xylem to mesophyll (hollow blue arrow), via the bundle sheath cells (BSCs), which tightly envelop the xylem. GC, stomatal guard cell; MC, mesophyll cell; PD, plasmodesma. (B) Blue-light (BL) signaling pathway (blue arrowheads) in a BSC, from BL perception by the phototropin receptors (phot) complexed with the kinases, BLUS1 and BHP (Hayashi et al., 2017; Inoue and Kinoshita, 2017), through a hypothesized intermediate tyrphostin 9 (TP9)-inhibitable phosphorylation (P) of an unknown substrate X by an unknown Raf-like kinase, down to the ultimate AHA2 activation by its phosphorylation (P). AHA2 activation results in proton extrusion via the pump, hyperpolarization (charge separation across the membrane: A^-^ / H^+^), cytosolic alkalinization (pH〈) and xylem sap acidification (pH®; also reported in Grunwald et al., 2021), presumably, at the expense of ATP breakdown to ADP with a transient phosphorylation (P) of the catalytic site on the pump protein, as expected from a P-type H^+^-ATPase. Xylem sap acidification enhances (black dotted arrow) water permeability (blue doted arrows) of the BSC’s plasma membrane. This is presumably due to enhanced permeability of aquaporins (AQP)s and underlies the increased P_f_ (osmotic water permeability) of the single BSC and, in turn, also the increased K_leaf_ (Grunwald et al., 2021). P_f_ in a *constant external pH* may be further increased by light (Fig. 8), due to, among other possibilities, cytosolic alkalinization (Fig. 5B), and/ or the AHA2-generated hyperpolarization (Fig. 5A), or, perhaps, even protein-protein interaction with the phots (these possibilities are not represented here) . Additional details can be found in the text. Water exiting the BSC via plasmodesmata between BSCs and the neighbouring MCs or via AQPs to the apoplast (blue dotted arrows and question marks) are included for completeness, although it is currently unknown to what extent these symplastic and apoplastic paths contribute to the radial water flow across the BSCs and whether they are also affected by BL

### An early K_leaf_ response to blue light - what is the advantage?

Both the opening of the stomata and the increase in K_leaf_ occur in the early morning hours (e.g., Brodribb and Holbrook, 2004; Domec et al., 2009; Locke and Ort, 2015) when the irradiance spectrum includes relatively high levels of BL (Matthews et al., 2020, and references therein). *What is the advantage of the early [blue-]light response of stomata?* We suggest that it allows the acquisition of CO_2_ under sub-maximum VPD (vapor pressure deficit, a measure of the driving force for leaf water evaporation), thereby increasing water-use efficiency.

Interestingly, K_leaf_ responds to light even faster than g_S_ does (Guyot et al., 2012) – in an ultimate manifestation of the independence of the BSCs’ BL signal transduction from that of the GCs’. One possible advantage of this almost-simultaneity is matching the stomatal opening with the increased water flux into the leaf. Were it not for this accompanying K_leaf_ increase, the BL-induced stomata opening, which peaks in the morning hours (the “golden hour”; (Gosa et al., 2019), could lead to an imbalance in leaf water, even in the presence of a relatively low VPD, leading to a drop in the leaf water potential (Ψ_leaf_), which, in turn, could close the stomata (Raschke, 1970; Guyot et al., 2012; Klein, 2014; Scoffoni et al., 2018) and thereby limit photosynthesis. Instead, an early morning increase in K_leaf_ elevates Ψ_leaf_, thereby increasing and prolonging stomatal opening (ibid). Hypothetically, under more extreme conditions (e.g., with high VPD even in the early morning), BL-increased K_leaf_ would prevent a hydraulic pathway failure when stomatal conductance is at its peak (Brodribb and Holbrook, 2004; Halperin et al., 2017; Gosa et al., 2019).

Another possible advantage of the early morning, BL-induced increase in K_leaf_ may be an increased CO_2_ permeability of aquaporins. A few lines of evidence converge to support this hypothesis; (a) CO_2_ can cross cell membranes through aquaporins, particularly aquaporins of the PIP1 family (Kaldenhoff and Fischer, 2006), (b) the transcripts of PIP1 family aquaporins in walnut have been shown to be upregulated concomitantly with the increase in K_leaf_ induced by a brief (15 min) exposure to white light (Ben Baaziz et al., 2012), (c) in soybean (*Glycine max*), the transcript of a PIP1 family aquaporin (PIP1;8) was found to be more abundant in the morning than at midday, in correlation with a higher K_leaf_ in the morning (Locke and Ort, 2015), (d) aquaporins control K_leaf_ (Shatil-Cohen and Moshelion, 2012; Sade et al., 2014), (e) CO_2_ transported by xylem from the roots may be a source for the CO_2_ assimilated into the bundle sheath and mesophyll (Janacek et al., 2009; Hubeau et al., 2019). Thus, assuming the K_leaf_ increase with the first light of day parallels the increase in the permeability of BSC aquaporins to CO_2_, the passage from the xylem to the mesophyll of the root-originated CO_2_ will be enhanced even before full stomatal opening. The increased availability of CO_2_ at this time, when the photosynthetic light is already sufficient, would enhance CO_2_ assimilation – a great advantage for the plant. This hypothesis is strengthened by the fact that (Stutz and Hanson, 2019) have shown that the xylem-transported CO_2_ constitutes a varying proportion of the total CO_2_ assimilated throughout the day depending on the availability of light and CO_2_. Notably, photosynthetic CO_2_ uptake reaches a maximum within 10 min of exposure to BL (Doi et al., 2015), which is over 3-fold faster than the full opening of stomata (Grondin et al., 2015).

### The advantage of diminished AHA activity and P_f_ at night

The fact that the BSC P_f_ is low in the dark (Figure 8) may explain how the mesophyll does not get flooded at night when root pressure increases. At night, the P_f_ and K_leaf_ reduction likely prevent water influx into the leaf, while the buildup of root pressure is relieved via guttation. A malfunction of this mechanism is perhaps what underlies leaf flooding by the xylem sap (Shatil-Cohen and Moshelion, 2012). The inactivity of the BSCs H^+^-ATPases, manifested here in the nightly alkalization of the xylem sap and in guttation drops, (as noted earlier, for example, also in tomatoes (Urrestarazu et al., 1996) and poplars (Aubrey et al., 2011) , could be advantageous for the plant as a means of energy conservation, as ATPases operate at a high energetic cost (Palmgren, 2001).

Our findings expand the understanding of the molecular basis of the regulation of leaf water influx. The fact that BL increases K_leaf_ in several other species imparts an even more general importance to our results. A focus on the hydraulic valve in series with the stomata should provide new directions for the study of plant water relations in different environments.

#### AUTHOR CONTRIBUTIONS

YG and SG constructed the BSCs-directed transgenic lines: mutant complementation and *PHOT* silencing., planned and performed the K_leaf_ and vein pH experiments, and wrote the paper. YG planned and performed the P_f_ experiments. TT planned and performed the membrane potential and cytosolic pH experiments. NM and MM conceptualized and supervised the project and wrote the paper.

## ACKNOWLEDGMENTS

We thank Dr. Dizza Bursztyn of the Hebrew University of Jerusalem for assisting us with the statistical analysis of this work. We thank Dr. Naama Gil and Dr. Idan Efroni for assisting in the RLM RACE *amiRNA* procedure.

This research was supported by the We thank Dr. Naama Gil and Dr. Idan Efroni for assisting in the RLM RACE *amiRNA* procedure Israel Science Foundation (ISF) grant No. 1043/20 to MM and the ISF grant No. 1312/12 to NM.

## SUPPLEMENTAL FIGURES AND TABLES

**Supplemental Figure S1.** The relationship between the di-8-ANEPPS fluorescence ratio (F-ratio, F_438_/F_531_) and the BSC membrane potential (E_M_).

**Supplemental Figure S2.** phot receptor mutants do not show any effect of BL on their g_S_, E or Ψ_leaf_.

**Supplemental Figure S3.** S*C*R*:amiR*-silencing of the *PHOT1* or *PHOT2* genes in the BSCs reduces Ψ_leaf_ under red + blue light, but does not reduce the stomatal conductance (g_S_) or the rate of transpiration (E).

**Supplemental Figure S4.** BSC-directed *PHOT1* complementation of the *phot1-5* mutant elevates the leaf water potential (Ψ_leaf_), but not stomatal conductance (g_S_) or the transpiration rate (E).

**Supplemental Figure S5.** Tyrphostin 9 perfused via the petiole does not affect the examined physiological parameters under red-light-only illumination.

**Supplemental Figure S6.** Xylem-fed kinase inhibitor tyrphostin 9 does not affect the blue light-induced increase in the transpiration rate (E), but does prevent any blue light-induced increase in leaf water potential (Ψ_leaf_).

**Supplemental Figure S7.** BSC-directed *AHA2* complementation of the *aha2-4* mutant restores its blue light-induced leaf water potential (Ψ_leaf_) increase, but not the blue light-induced increases in stomatal conductance (g_S_) or in the transpiration rate (E).

**Supplemental Figure S8.** Leaf vein density does not depend on the expression of phot1 or phot2.

**Supplemental Figure S9.** After prolonged illumination, stomatal conductance (g_S_) of intact leaves of whole plants did not differ among the different genotypes (WT, *phot* mutants and *SCR:PHOT1*).

**Supplemental Figure S10.** The BLUS1 kinase mutation abolishes the BL-induced K_leaf_ increase.

**Supplemental Figure S11.** The hyperpolarization of BSCprotoplasts by BL is prevented by the phot1 mutation.

**Supplemental Table S1.** List of primers used in this study

**Supplemental Table S2.** Expression levels of some genes found in the transcriptomes of bundle-sheath (BS) cells and mesophyll (MES) cells (Wigoda et al., 2017, GEO repository Experiment GSE85463) that are or may be related to blue-light signaling, encoding receptors, kinases, H^+^-ATPases and aquaporins.

## Notes

### Competing Interest Statement

The authors have declared no competing interest.

### Summary of Updates

All figures were converted from "bar graphs" to "box and whiskers"

