## Supplemental Figures1-11 and tables 1-2 for "Out of the blue: Phototropins of the leaf vascular bundle sheath mediate the regulation of leaf hydraulic conductance by blue light"

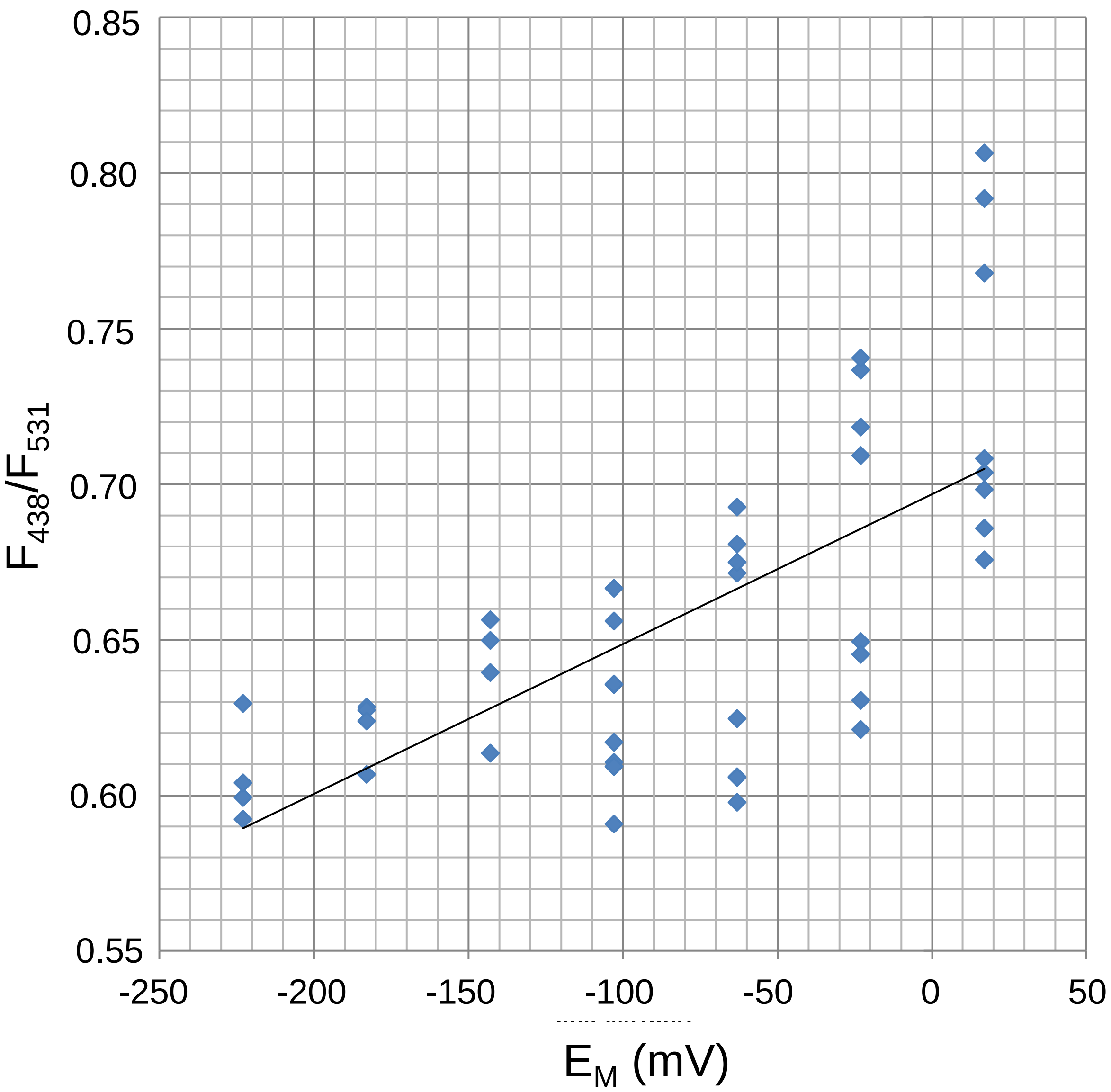


**Supplemental Figure S1.** The relationship between the di-8-ANEPPS fluorescence ratio (F-ratio, $F_{438}/F_{531}$) and the BSC's membrane potential ($E_M$). The F-ratio was obtained from a pair of images recorded from a BSC protoplast at the indicated excitation wavelengths while its membrane potential ($E_M$) was imposed using a patch-clamp pipette (Materials and methods). The linear fit to the data points was: F-ratio = 0.0005*$E_M$ + 0.6967 ($R^2$ = 0.4596). Note the positive correlation between the F-ratio and $E_M$ ($P<4.3\ 10^{-7}$). *Supports Figure 5A.*

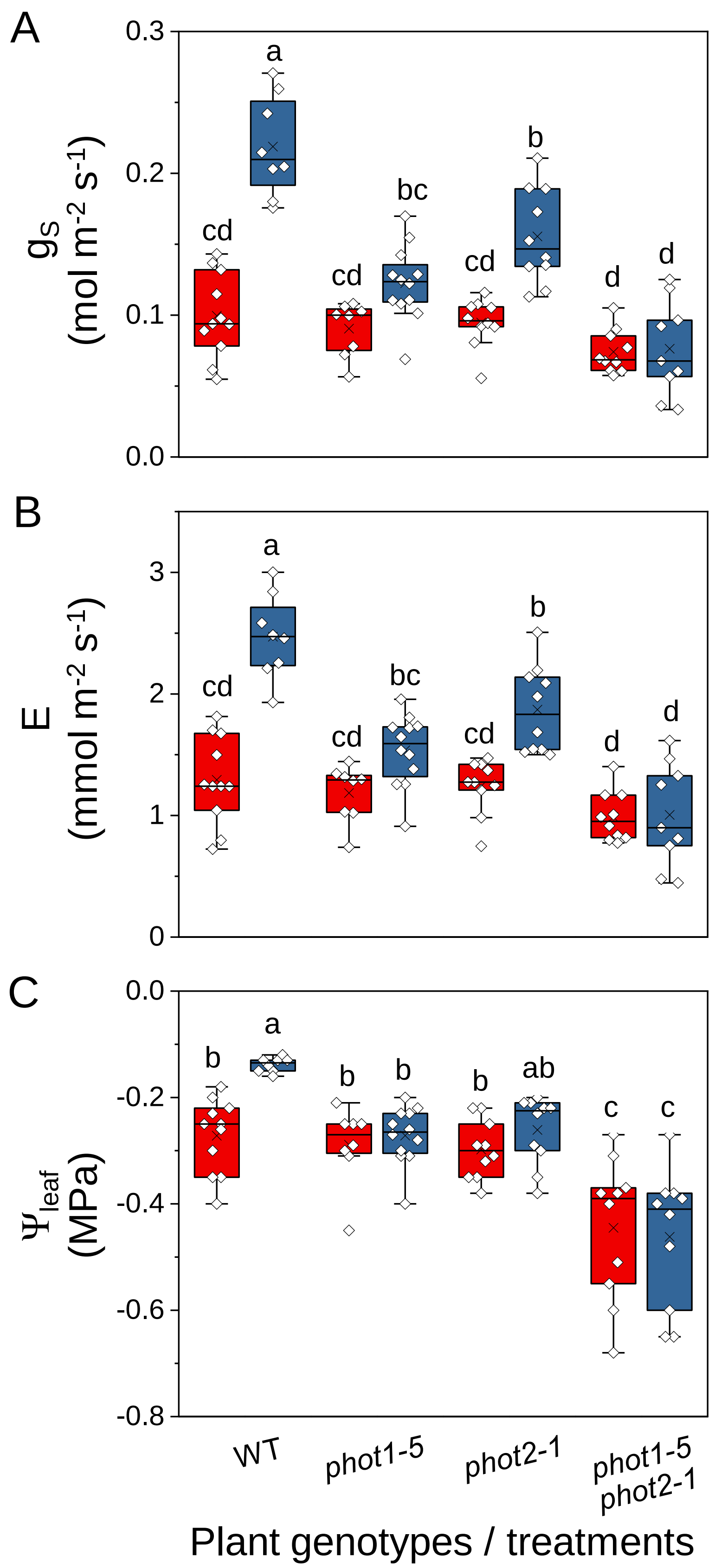


**Supplemental Figure S2.** phot receptor mutants do not show any effect of BL on their $g_S$, E or $\Psi_{leaf}$. (A) Stomatal conductance ($g_S$), (B) transpiration rate (E) and (C) leaf water potential **(**$\Psi_{leaf}$) of fully expanded detached leaves of WT (ecotype gl) and *PHOT* mutants (*phot1-5*, *phot2-1* and a double mutant, *phot1-5 phot2-1*) after illumination for 15 min immediately after dark, with RL (220 µmol m$^{-2}$ s$^{-1}$) or RL+BL (220 µmol m$^{-2}$ s$^{-1}$ with 90% RL and 10% BL). The box plot details are as in Fig. 1. Different letters denote significantly different values (ANOVA, post hoc Tukey's test; $P < 0.05$). Note the diminished response to RL+BL in the mutant lines. *Supports Figure 1 (via Eq. 1).*

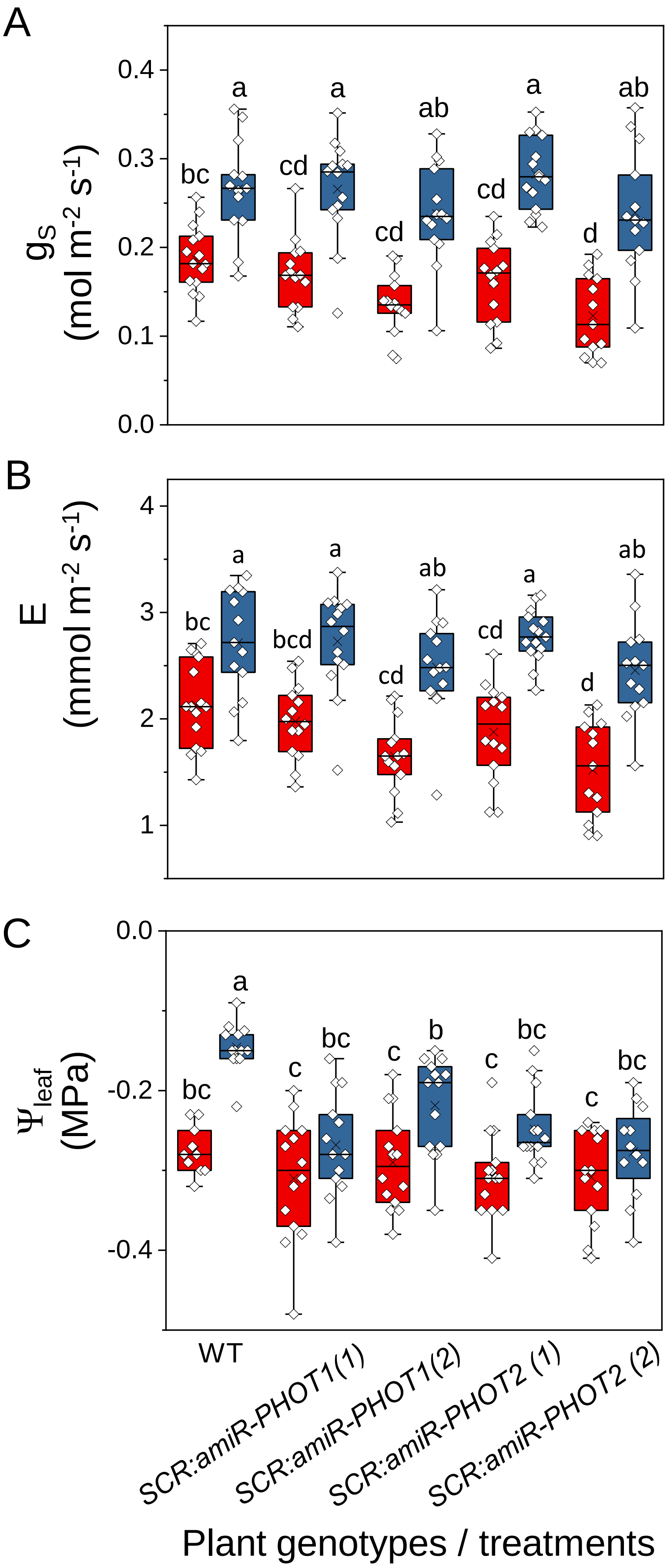


**Supplemental Figure S3.** *SCR:amiR*-silencing of the *PHOT1* or *PHOT2* genes in the BSCs reduces $\Psi_{leaf}$ under RL+BL, but does not reduce stomatal conductance ($g_S$) or the rate of transpiration (E). (A) $g_S$, (B) E and (C) leaf water potential ($\Psi_{leaf}$) of *PHOT1*- and *PHOT2*-silenced (*SCR:amiR-phot*) plants. *SCR:amiR-phot* plants were generated in the WT (Col-0) background. Fully expanded leaves of WT, *SCR:amiR-phot1* and *SCR:amiR-phot2* plants were treated with light as described in Figure 1. Other details also as in Fig. 1. Note that in plants with *PHOTs amiR*-silenced solely in their BSCs, $g_S$ and E remained unaffected under the RL+BL regime, while $K_{leaf}$ was impaired (Figure 2). *Supports Figure 2 (via Eq. 1).*

Grunwald, Gosa et al., 2022

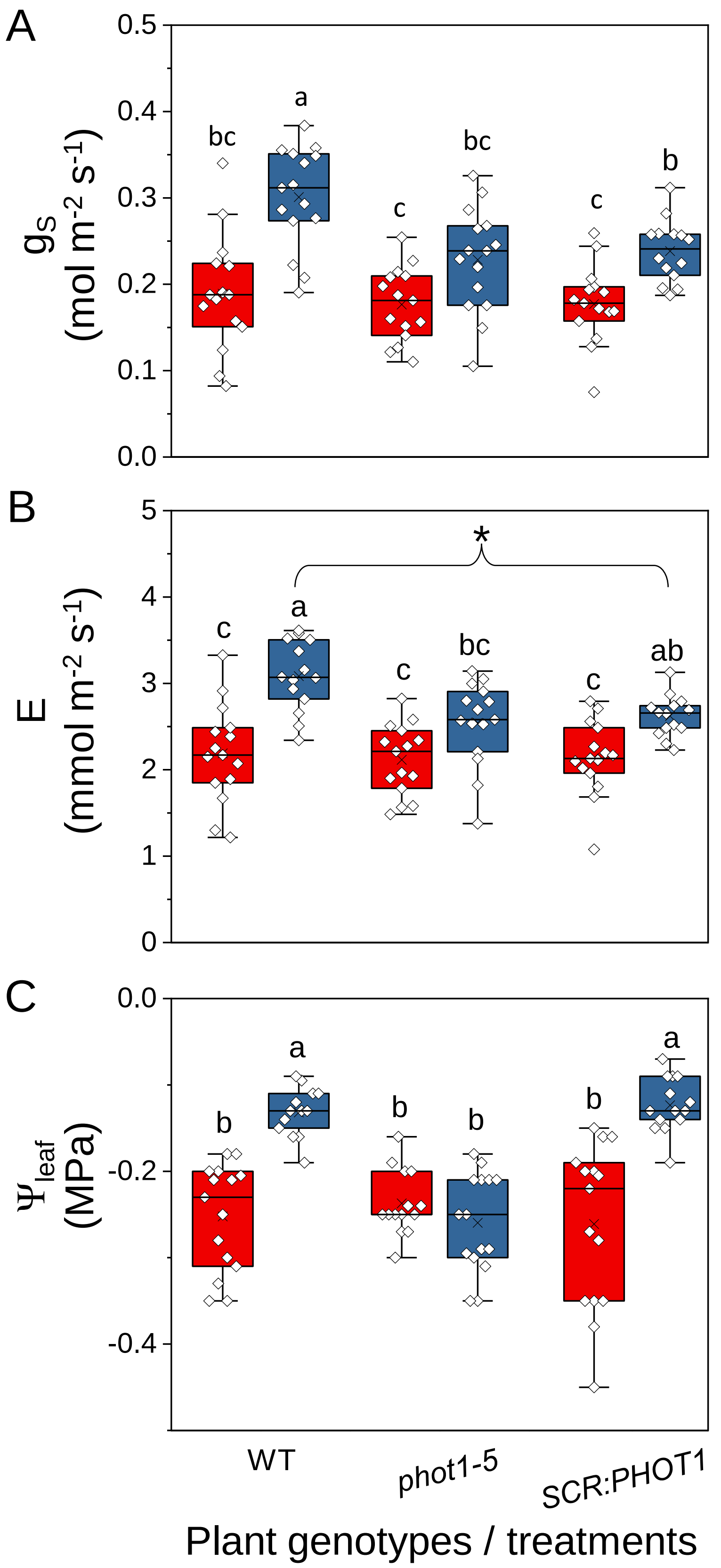


**Supplemental Figure S4.** BSC-directed *PHOT1* complementation of the *phot1-5* mutant elevates the leaf water potential ($\Psi_{leaf}$), but not the stomatal conductance ($g_S$) or the rate of transpiration (E). (A) $g_S$, (B) E and (C) $\Psi_{leaf}$. *phot1-5* (Col-gl) plants were complemented with *SCR*:*PHOT1* restoring phot1 activity specifically in the BSCs. Fully expanded leaves of WT (Col-gl), *phot1-5* and *SCR*:*PHOT1* were illuminated as described in Figure 1. Other details are also as in Fig. 1. Note the significant difference between the $g_S$ values of WT and *SCR*:*PHOT1* under RL+BL. The asterisk indicates a significant difference between the E values of WT and *SCR*:*PHOT1* under RL+BL (at *P* = 0.05, by a two-tailed, unpaired, equal variance *t*-test). *Supports Figure 3 (via Eq. 1).*

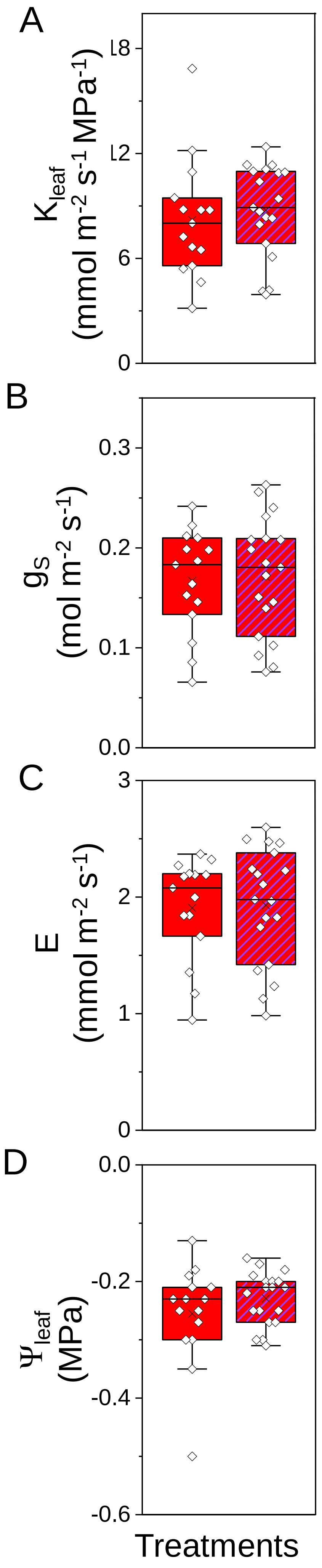


**Supplemental Figure S5**. Tyrphostin 9 perfused via the petiole does not affect the examined physiological parameters under RL-only illumination. Fully expanded leaves of WT (Col-gl) were pre-incubated (petiole deep) in AXS without (plain boxes) or with tyrphostin 9 (10 µM; hatched boxes) and kept overnight in dark boxes. Immediately after dark, leaves were illuminated for 15 min with RL (220 µmol $m^{-2}$ $s^{-1}$). (A) $K_{leaf}$, (B) $g_S$, (C) E, and (D) $\Psi_{leaf}$. Other details as in Fig. 4. *Supports Figure 4 as a control.*

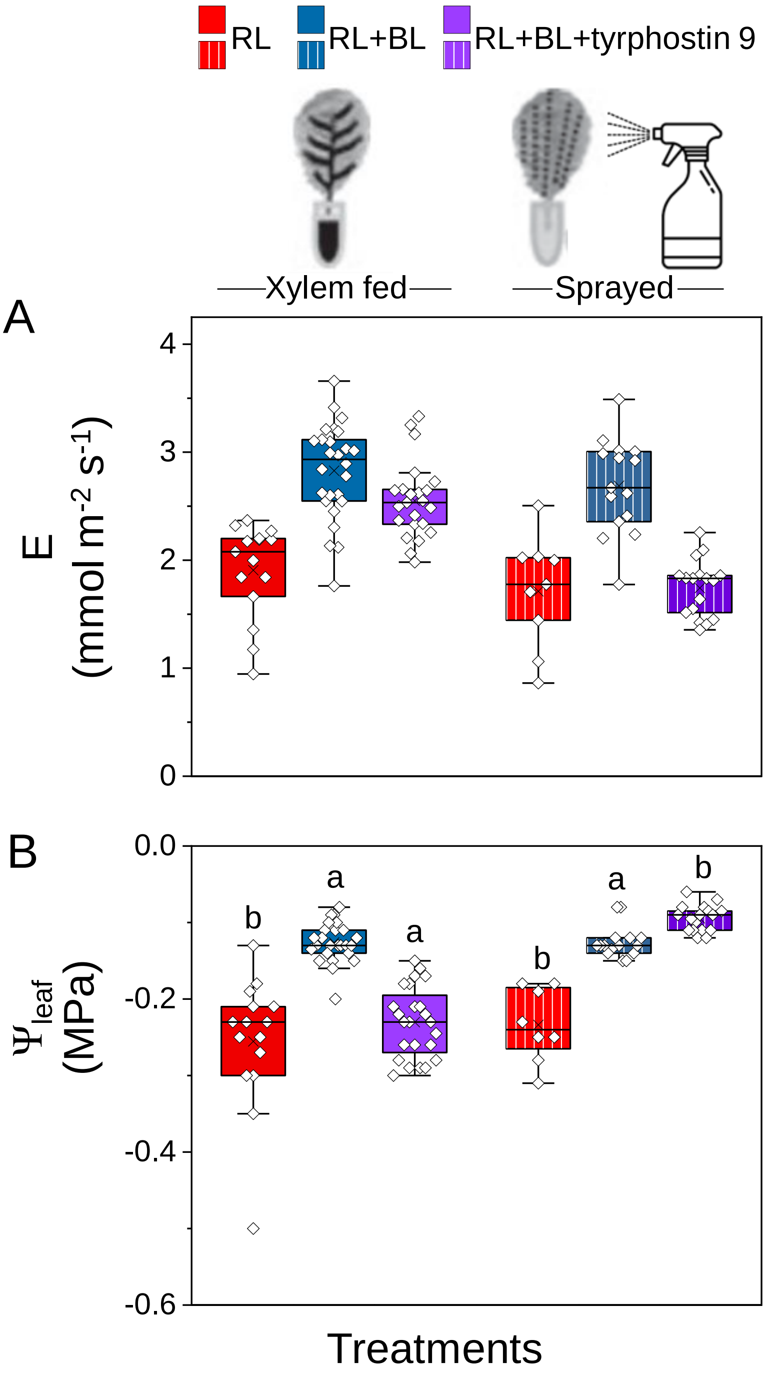


**Supplemental Figure S6**. Xylem-fed kinase inhibitor, tyrphostin 9, does not affect the BL-induced increase in the transpiration rate (E), but does prevent any BL-induced increase in leaf water potential ($\Psi_{leaf}$). Tyrphostin 9 sprayed on leaves acts in the opposite manner. (A) E and (B) $\Psi_{leaf}$. Different letters denote significantly different values (ANOVA, post hoc Tukey's test; $P < 0.05$ for all six treatments). Other details as in Fig. 4. *Supports Figure 4 (via Eq. 1).*

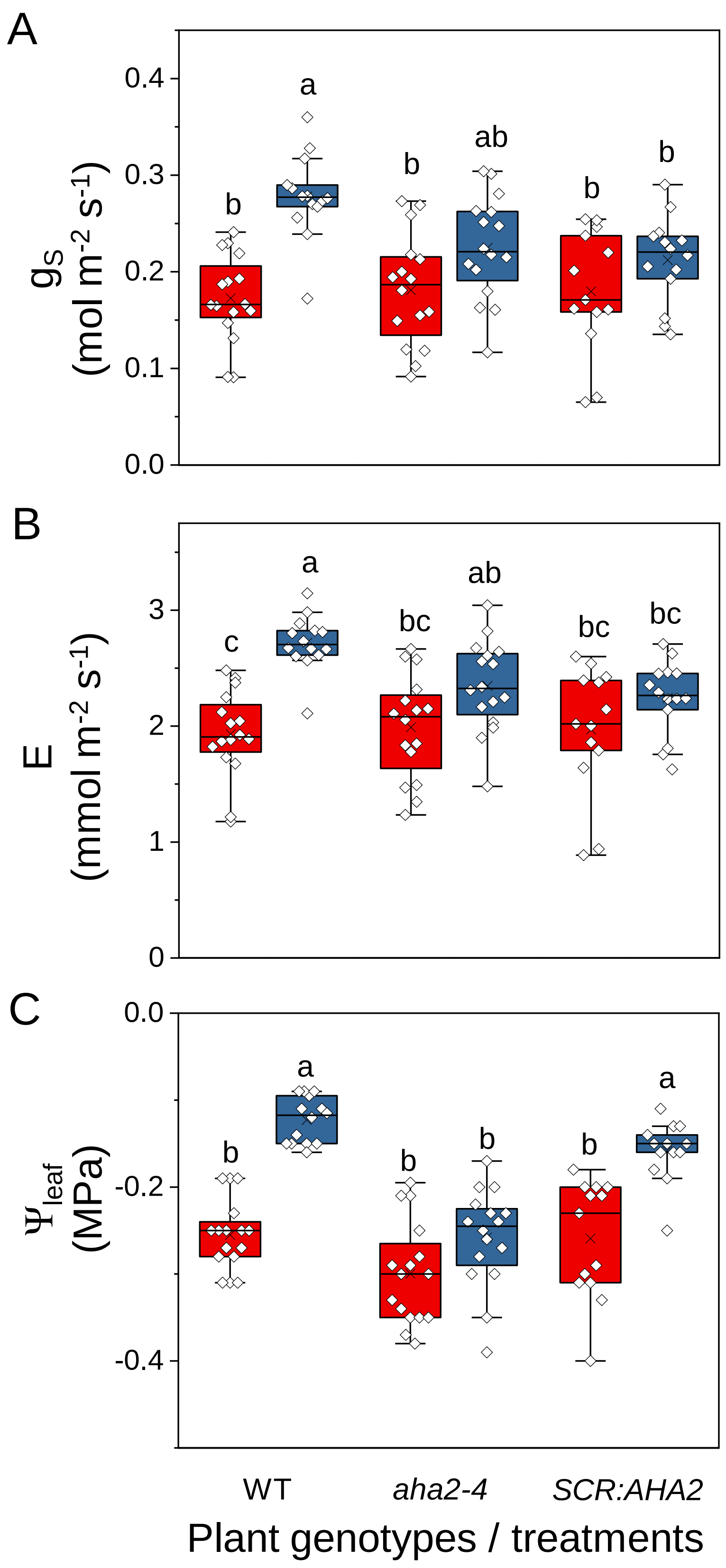


**Supplemental Figure S7.** BSC-directed *AHA2* complementation of the *aha2-4* mutant restores its BL-induced leaf water potential ($\Psi_{leaf}$) increase, but not the BL-induced increases in stomatal conductance ($g_S$) or in the transpiration rate (E). (A) $g_S$, (B) E and (C) $\Psi_{leaf}$ of fully expanded detached leaves of WT (Col-0) plants, *aha2-4* (Col-0) mutant plants and *aha2-4* plants complemented solely in their BSCs with *SCR:AHA2,* illuminated for 15 min with RL or RL+BL, as in Figure 1. Other details also as in Figure 1. *Supports Figure 6 (via Eq. 1).*

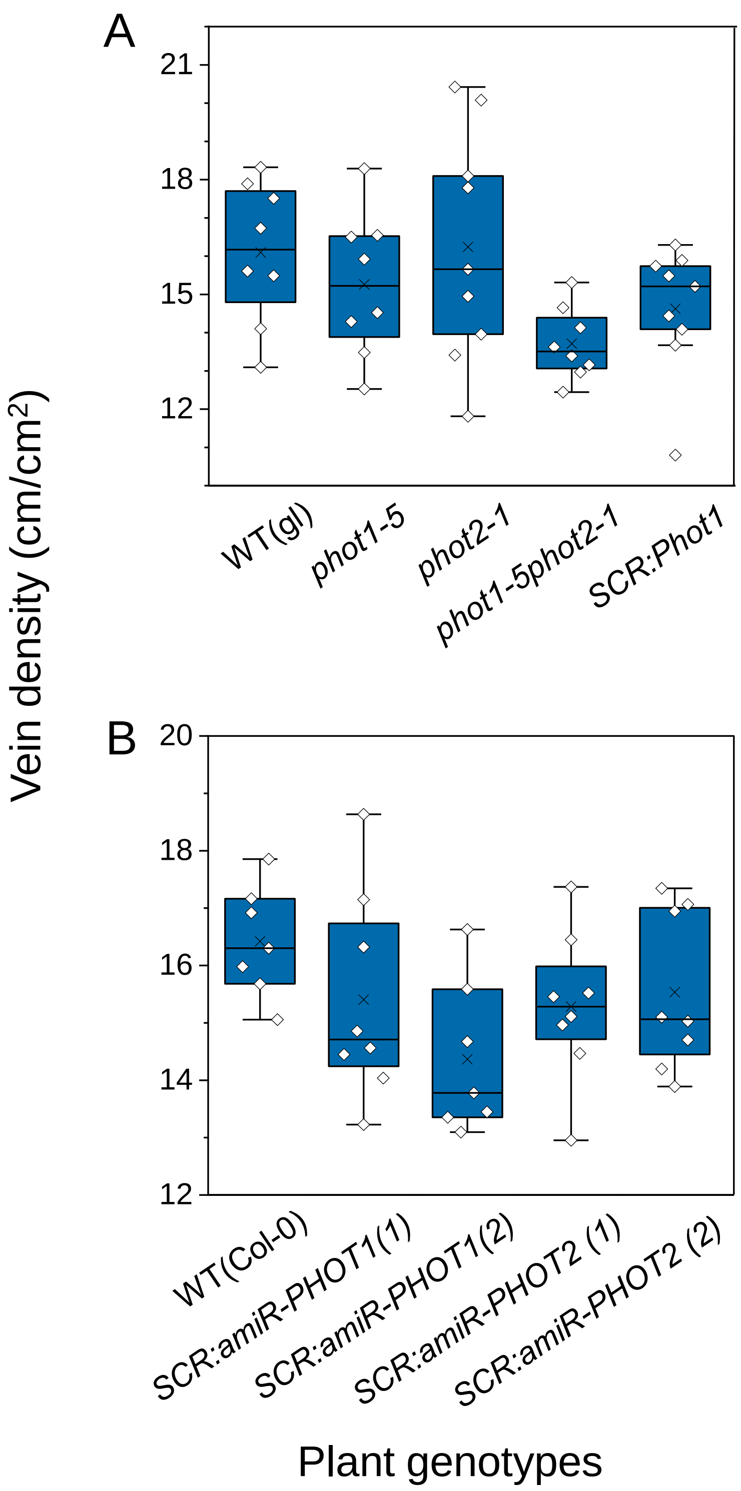


**Supplemental Figure S8**. Leaf vein density does not depend on the expression of phot1 or phot2. Vein density averaged over the indicated number of leaves). Vein density was calculated as the total vein length divided by the scanned leaf area. (A) phot mutants compared to the WT. (B) *PHOT*-silenced lines compared to the WT. Statistics as in Fig 1. Note the lack of any significant differences between the different mutants and *PHOT*-silenced lines and their WT controls. *Supports Figure 2, as controls.*

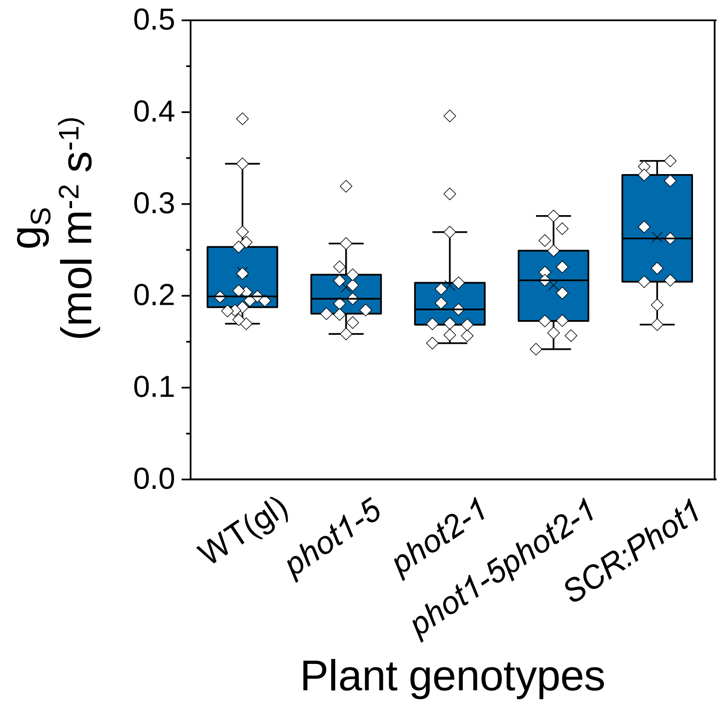


**Supplemental Figure S9.** After prolonged illumination, stomatal conductance ($g_S$) of intact leaves of whole plants did not differ among the different genotypes (WT, *phot* mutants and *phot1-5* complemented with *SCR:PHOT1*).The plants were left under the growth-room lights for 2–4 h after the lights were turned on. Statistics – as in Fig 1. *Supports Figures 1 and 3 as controls*.

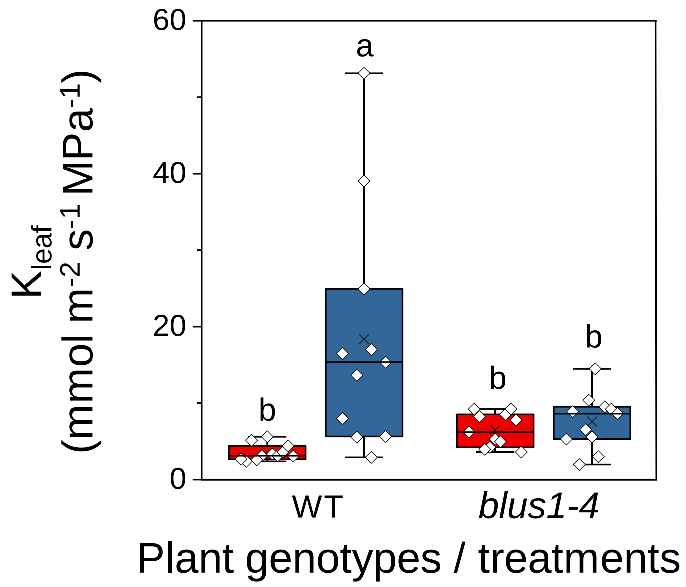


**Supplemental Figure S10.** The BLUS1 kinase mutation abolishes the BL-induced $K_{leaf}$ increase. $K_{leaf}$ were determined in fully expanded leaves of WT (Col-0) and BLUS1 kinase mutant *blus1-4* (Col-0) plants immediately after light treatments as described in Figure 1 legend. Note the absence of the response to RL+BL in the mutant line. Statistics – as in Fig 1. *Supports Figures 1-3 in establishing the similarity of the BSCs' BL-signaling initiation to that in GCs.*

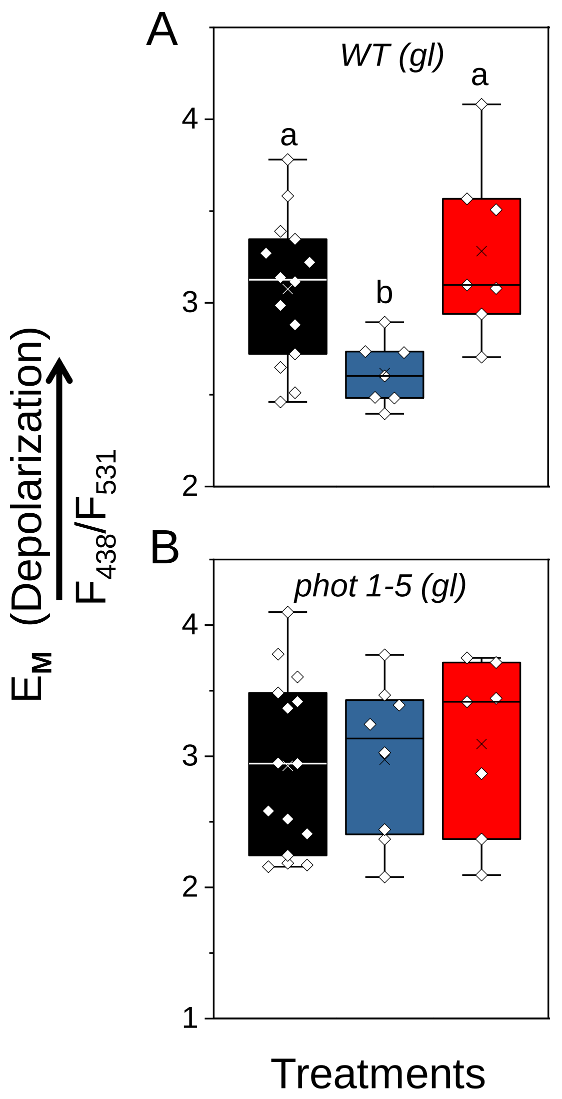


**Supplemental Figure S11.** The hyperpolarization of BSC protoplasts by BL is prevented by the mutation of phot1. Membrane potential ($E_M$, in units of fluorescence ratio) of protoplasts from WT and mutant plants without the *SCR:*GFP label, with a probable (at least 70 % chance) BSC identity (Materials and Methods) was assayed under dark, RL or RL+BL illumination. $E_M$ was reported by the dual-excitation, ratiometric fluorescent dye di-8-ANEPPS. (A) WT (Col-gl) protoplasts exposed to RL+BL were hyperpolarized relative to those in the dark or under RL alone. (B) *phot1-5* (Col-gl), the phot1 mutant, failed to respond to RL+BL. Other details are as in Fig. 5A, except the range of the F-ratio values, due to an update of the excitation light source. Importantly, a new calibration that we performed, like in Suppl. Fig. S1, also revealed a positive correlation between the fluorescence ratio and the membrane potential, which enables a comparison between Fig. 5A and S11 with respect to the *direction* of the $E_M$ change (depolarization or hyperpolarization).

**Supplemental Table S1**. List of primers used in this study

| **AHA 2 Genotyping** | **5' -> 3' Primer Sequence** |
| --- | --- |
| *aha2-4* forward | ATGTTCATTGCAAAGGTGGTG |
| *aha2-4* reverse | CCCATTAGCTCGTGGTTATTG |
| LBb1.3 tDNA insert | ATTTTGCCGATTTCGGAAC |
| T-DNA LB | TCAAACAGGATTTTCGCCTGCT |
| **AHA2 Trans. Genotyping** | |
| *aha2-4* forward 2972 | ATGATTGCTTTCCTCATTGCA |
| 35S terminator reverse | GCAGGTCACTGGATTTTGGT |
| gateway ends for cloning | |
| B1F | GGGGACAAGTTTGTACAAAAAAGC |
| B2R | GGGGACCACTTTGTACAAGAAAG |
| ***SCR:amiR-PHOT*** **sequencing** | |
| *SCR*-2341F | TTCTCCTCATCGGAGATCGT |
| *amiR*_Seq_F | GCTCGGACGCATATTACACA |
| *amiR*_Seq_R | CCAATTTGTCTACCGCATCA |
| ***PHOT1*** **cDNA amplification for cloning** | |
| *PHOT1*kin_F | CAGAGTGGGGATTGGTTTTGAAG |
| *PHOT1*kin_R | GTTTGCTGATCTTCTAGCTCAGG |

**Supplemental Table S2**. Expression levels of some genes found in the transcriptomes of bundle-sheath (BS) cells and mesophyll (MES) cells (Wigoda et al., 2017, GEO repository Experiment GSE85463) that are or may be related to blue-light signaling, encoding receptors, kinases, $H^+$-ATPases and aquaporins. Mean $\log_2$ normalized raw expression levels are presented with their standard errors (SE). The choice of kinases was based on Hayashi et al. (2017).

| Gene Title | AGI | $Log_2$ (Mean_BS) | $Log_2$ (Mean_MES) | SE_BS | SE_MES |
|---|---|---|---|---|---|
| **Blue-light receptors** | | | | | |
| PHOT1 (phototropin 1) | AT3G45780 | 1.70 | 1.75 | 0.06 | 0.02 |
| PHOT2 (phototropin 2) | AT5G58140 | 3.31 | 3.48 | 0.20 | 0.14 |
| **Phot-interacting kinase** | | | | | |
| BLUS1 (blue light signaling1) | AT4G14480 | 1.77 | 1.71 | 0.13 | 0.03 |
| **Raf-like kinases (blue light downstream signaling genes), based on Hayashi et al. (2017)** | | | | | |
| BHP (blue light-dependent H+-ATPase phosphorylation) | AT4G18950 | 7.36 | 7.04 | 1.24 | 0.68 |
| VIK (VH1-interacting kinase) | AT1G14000 | 5.49 | 5.05 | 0.35 | 0.28 |
| **Plasma membrane $H^+$-ATPases** | | | | | |
| AHA1 (Arabidopsis $H^+$-ATPase 1) | AT2G18960 | 5.71 | 4.60 | 1.16 | 0.97 |
| AHA2 (Arabidopsis $H^+$-ATPase 2) | AT4G30190 | 5.46 | 3.81 | 0.74 | 0.27 |
| AHA3 (Arabidopsis $H^+$-ATPase 3) | AT5G57350 | 1.70 | 1.62 | 0.10 | 0.05 |
| AHA4 (Arabidopsis $H^+$-ATPase 4) | AT3G47950 | 1.86 | 1.87 | 0.11 | 0.12 |
| AHA5 (Arabidopsis $H^+$-ATPase 5) | AT2G24520 | 1.66 | 1.73 | 0.02 | 0.16 |
| AHA6 (Arabidopsis $H^+$-ATPase 6) | AT2G07560 | 2.08 | 2.27 | 0.09 | 0.03 |
| AHA7 (Arabidopsis $H^+$-ATPase 7) | AT3G60330 | 1.61 | 1.80 | 0.06 | 0.06 |
| AHA8 (Arabidopsis $H^+$-ATPase 8) | AT3G42640 | 2.22 | 2.19 | 0.03 | 0.08 |
| AHA9 (Arabidopsis $H^+$-ATPase 9) | AT1G80660 | 1.91 | 1.49 | 0.34 | 0.09 |
| AHA10 (auto-inhibited $H^+$-ATPase isoform 10) | AT1G17260 | 1.67 | 1.63 | 0.02 | 0.01 |
| AHA11 (Arabidopsis $H^+$-ATPase 11) | AT5G62670 | 3.59 | 4.38 | 0.52 | 0.95 |
| **Aquaporins** | | | | | |
| PIP1;1 (Plasma membrane intrinsic protein 1;1) | AT3G61430 | 3.25 | 2.40 | 0.71 | 0.11 |
| PIP1;2 (Plasma membrane intrinsic protein 1;2) | AT2G45960 | 2.79 | 2.15 | 0.72 | 0.42 |
| PIP1;3 (Plasma membrane intrinsic protein 1;3) | AT1G01620 | 2.47 | 2.50 | 0.14 | 0.20 |
| PIP1;4 (Plasma membrane intrinsic protein 1;4) | AT4G00430 | 4.76 | 5.02 | 0.38 | 0.53 |
| PIP1;5 (Plasma membrane intrinsic protein 1;5) | AT4G23400 | 2.27 | 2.36 | 0.18 | 0.35 |
| PIP2;1 (Plasma membrane intrinsic protein 2;1) | AT3G53420 | 3.67 | 3.63 | 0.66 | 0.78 |
| PIP2;4 (Plasma membrane intrinsic protein 2;4) | AT5G60660 | 2.06 | 2.12 | 0.02 | 0.02 |
| PIP2;5 (Plasma membrane intrinsic protein 2;5) | AT3G54820 | 2.44 | 2.48 | 0.06 | 0.07 |
| PIP2;6 (Plasma membrane intrinsic protein 2;6) | AT2G39010 | 4.98 | 4.32 | 1.01 | 0.42 |
| PIP2;7 (Plasma membrane intrinsic protein 2:7) | AT4G35100 | 4.94 | 4.10 | 1.14 | 0.72 |
| PIP2;8 (Plasma membrane intrinsic protein 2;8) | AT2G16850 | 1.77 | 1.61 | 0.04 | 0.12 |
